# Evolutionary origins and chromatin state shape X-chromosome upregulation pattern during eutherian and metatherian embryogenesis

**DOI:** 10.64898/2026.04.24.720575

**Authors:** Hemant C Naik, Pavithra Narendran, Srimonta Gayen

## Abstract

In mammals, silent state of one of the X-chromosomes in female balance the X-dosage between sexes. In parallel, X to autosome imbalance due to monoallelic expression of X-linked genes relative to biallelic autosomal genes, is primarily compensated through the X-chromosome upregulation (XCU). It has been demonstrated that X-chromosome inactivation (XCI) and XCU coincides during embryogenesis, however, XCU is not global and it occurs in a gene-specific manner. The underlying mechanistic aspects of such specificity of XCU remain unknown. Here, we provide systematic and comparative analysis across eutherians (mouse, human and pig) and metatherian (opossum) embryogenesis to determine if evolutionary origins shape the XCU. Intriguingly, we show that while evolutionary older X-linked genes (predating mammalian divergence) undergo robust XCU consistently across developmental stages, younger mammalian origin genes do not. Similarly, the eutherian X-linked genes conserved in metatherian X (X-ortholog) undergo robust XCU, whereas genes orthologus to metatherian autosome (Auto-ortholog) exhibit weaker pattern of XCU. Further, strata-wise comparison revealed that genes in older XLJchromosome strata (1–2) consistently undergo upregulation, whereas strata 3–4 genes do not. Importantly, we show that different evolutionary classes of X-linked genes, which undergo robust XCU, are often enriched with active chromatin marks (H3K36me3, H3K27ac and H3K4me1) relative to the autosome, suggesting that chromatin state mediate the XCU. Moreover, we show that often active-marks enrichment correlates with differential XCU dynamics of different class of genes. Taken together, our study provides significant insight into the evolutionary dynamics of XCU and underlying mechanistic framework.

## Introduction

In mammals, inactivation of one X-chromosome in female cells compensates the dosage of X-linked gene expression between sexes, resulting in monoallelic expression from X (Lyon 1961). However, this creates a transcriptome-wide dosage inequality of X-linked genes with the biallelic autosomal genes similar to male cells. To compensate for this dosage deficit, majority of X-linked genes on the active X chromosome are upregulated in both males and females, a process known as X-chromosome upregulation (XCU) (Ohno S 1967; Di and Disteche 2006; Gupta et al. 2006; Lin et al. 2007, 2011; Deng et al. 2011; Sangrithi et al. 2017; Cecalev et al. 2024; Graves 2016). Recent studies using allele-specific RNA sequencing have disentangled expression patterns of X-linked genes from active versus inactive X chromosomes and demonstrated that XCU is coordinated with the silencing of X-linked genes on the inactive X in early mouse embryos (Lentini et al. 2022; Naik et al. 2022; Wei et al. 2024). Moreover, loss of one X chromosome in mouse embryonic stem cells (mESC) or induced pluripotent stem cells (iPSCs) leads to upregulation of the X-linked genes from the other X (Talon et al. 2021; Lentini et al. 2022). Interestingly, reactivation of the inactive-X in female cells triggers the erasure of XCU in epiblast cells and during the reprogramming of fibroblast to iPSCs (Naik et al. 2024; Lentini et al. 2022; Talon et al. 2021). Together, the XCU are tightly linked to the inactivation or loss of the other X-chromosome in female cells. Although, XCU occurs in a coordinated fashion with the XCI, it has been shown that XCU is not global rather it happens in a gene-by-gene basis (Naik et al. 2022). Additionally, it was shown that transposable elements do not undergo XCU (Wei et al. 2024). Recently, it has also been shown that heterozygous deletion of some parts of the X-chromosome in mESC leads to upregulation of genes from the counterparts, but not for all genes (Allsop et al. 2025). However, underlying basis of gene-specific XCU remains poorly understood.

It has been shown that often evolutionary age of genes is connected to dosage compensation behavior. It was shown that kinetics of XCI of evolutionary older vs. younger genes differs during imprinted XCI in mouse (Wei et al. 2024). Moreover, XCI pattern and mechanisms involved often varies across species (Alfeghaly and Rougeulle 2025; Grant et al. 2012). Importantly, it has been demonstrated that the frequency of genes that escape X-inactivation varies across eutherian and metatherian species (Balaton et al. 2021; Wang et al. 2014; Rodríguez-Delgado et al. 2014; Al Nadaf et al. 2012). Similarly, it has been shown that pattern of XCU differs between eutherian and metatherian (Zhang et al. 2025; Julien et al. 2012). In this study, we have systematically explored whether evolutionary origins of X-linked genes govern the XCU dynamics during early embryogenesis across different mammalian species: eutherian (mouse, human, pig) and metatherian (opossum). For this, we have classified X-linked genes into different evolutionary subgroups: (1) genes predated before mammalian origin (old) vs. mammalian-origin genes (young); (2) X-linked genes conserved between eutherian and metatherian X (X-ortholog) vs. X-linked genes orthologous to marsupial autosome (Auto-ortholog); (3) genes belonging to different evolutionary strata (strata1, strata2 and strata3+4) (Lahn and Page 1999; Pandey et al. 2013; Sandstedt and Tucker 2004).

On the other hand, molecular factors and mechanism governing XCU remains underexplored. It has been demonstrated that enhanced chromatin accessibility is associated with the XCU (Talon et al. 2021). However, the role of different chromatin modifiers in driving XCU remains unknown. While few studies indicated that enrichment of different active histone modifications are linked to the XCU, other studies do not find much correlation between them (Lentini et al. 2022; Yildirim et al. 2012; Deng et al. 2013; Naik et al. 2022; Allsop et al. 2025). Here, we have explored the role of different active histone modifications in driving XCU of different evolutionary classes of genes during the early embryogenesis.

## Results

### Young X-linked genes undergo active-X upregulation at post-implantation, not pre-implantation, during imprinted XCI in mouse

In mouse, upon zygotic genome activation, the paternal X-chromosome in female cells undergoes inactivation, a phenomenon known as imprinted XCI (Huynh and Lee 2003; Okamoto et al. 2004; Takagi and Sasaki 1975). Subsequently, in pre-gastrulation epiblast, another round XCI occurs, where either the paternal or maternal X is randomly inactivated specified as random XCI (Takagi and Sasaki 1975; Saiba et al. 2018; Gayen et al. 2016; Samanta et al. 2022). Switching of imprinted to random happens through the reactivation of the inactive-X at the early epiblast cells of late blastocyst (Mak et al. 2004; Okamoto et al. 2004). In this study, to explore the active-X upregulation dynamics of genes with different evolutionary origins during the initiation of imprinted and random XCI in mouse, we have used available single-cell RNA-sequencing (scRNA-seq) datasets of hybrid mouse pre- and post-implantation embryos for our analysis (Fig. S1A) (Deng et al. 2014; Borensztein et al. 2017b, 2017a; Chen et al. 2016; Cheng et al. 2019). Additionally, we have used scRNA-seq datasets of hybrid mouse embryonic stem cells (mESCs) differentiation, which conventionally used as *ex vivo* model to study the initiation of random XCI (Fig. S1A) (Lentini et al. 2022; Böttcher et al. 2020). Since these embryos and mESCs were derived from the cross of evolutionary diverged mouse strains, they had single nucleotide polymorphisms (SNPs) between paternal vs. maternal genomes (Fig. S1A). Using these SNPs, we disentangled the expression of genes originating from maternal vs. paternal alleles through allele-specific analysis of scRNA-seq data. First, we segregated cells based on their XCI status as these cells are in the process of initiation and establishment of imprinted or random XCI. We classified cells of pre-and post-implantation embryos presumably into three categories based on their XCI status through analyzing the fraction allelic expression of the X-linked genes from the maternal-X: (1) XaXa: cells with no XCI (fraction X^mat^ expression: 0.4-0.6) (2) XaXp: cells with partial inactive-X (fraction X^mat^ expression: 0.6-0.8/0.2-0.4) (3) XaXi: cells with robust X-inactivation (fraction X^mat^ expression: 0.8-1/ 0.0-0.2) (Fig. S1B). On the other hand, autosomal genes showed equivalent allelic expression from both maternal and paternal allele, thereby validating our allele-specific expression quantification method (Fig. S1B). Moreover, as expected, the expression of X-linked genes in male cells were originated from maternal allele only in our analysis (Fig. S1B). Furthermore, cells from embryos having *Xist* deletion on the paternal-X (ΔXist-Pat), which do not undergo XCI (Borensztein et al. 2017b), showed equivalent allelic expression of X-linked genes from maternal and paternal X-chromosomes (Fig. S1B). Similar way we segregated XaXa, XaXp and XaXi cells in differentiated mESCs through calculating the fraction allelic expression of the X-linked genes from the C57 allele (Fig. S1C and S1D).

Next, we explored the dynamics of upregulation of active-X genes based on their evolutionary age: genes predated before mammalian origin (old) vs. mammalian-origin genes (young) (Fig. 1A). To delineate the active-X upregulation dynamics, we profiled X to autosome expression ratio (X:A) in allele-specific manner. If X-linked genes are not upregulated the expected allelic active-X:A ratio should be close to 1, whereas the allelic X:A ratio for upregulated X-linked genes are expected to be >1 and close to 2. We excluded low expressed genes (<1 TPM) and genes with very high expression (above 98 percentile) from our analysis. Surprisingly, we find that the ratio of paternal X (X^pat^): paternal-A (A^pat^) and maternal X (X^mat^): maternal-A (A^mat^) in XaXa cells (WT and ΔXist-Pat) of pre-implantation embryos for old genes was above 1 and close to 1.5, indicating moderate upregulation of both of the X-chromosomes (Fig. 1B). However, we did not observe such trend for young genes (Fig. 1B). Next, we compared the allelic X:A ratio of old (old-X:old-Auto) and young genes (young-X:young-Auto) between XaXa (WT and ΔXist-Pat) vs. XaXi cells of pre-implantation embryos (Fig. 1B). We observed that paternal X (X^pat^): A ratio of both old and young genes significantly decreased in XaXi cells compared to the XaXa cells and reached close to zero, indicating both old and young genes undergo imprinted XCI in pre-implantation embryos (Fig. 1B). Post-implantation visceral endoderm (VE) and extraembryonic ectoderm (ExE) also showed robust inactivation of old and young X-linked genes (Fig. 1B). On the other hand, active-X^mat^:A^mat^ ratio of old genes in XaXi cells was significantly increased compared to the XaXa cells in pre-implantation embryos as well as post-implantation VE and ExE, indicating upregulation of old X-linked genes from the active-X upon XCI (Fig. 1B). However, we did not observe such trends in case of young X-linked genes in pre-implantation embryos (Fig. 1B). But, we observed significant increase of active-X^mat^:A^mat^ ratio of young genes in post-implantation ExE and VE lineages, suggesting that young X-linked genes are upregulated in post-implantation embryos, not in pre-implantation. We excluded XaXp cells as they can represent cells of both classes: cell undergoing XCI or undergoing reactivation of the inactive-X. Further we validated our observation through comparing the active-X^mat^ expression with the A^mat^ expression in XaXi cells. Indeed, we found that while the expression of old X-linked genes from X^mat^ were significantly higher compared to the A^mat^ genes, there was no significant differences of young genes between X^mat^ vs. A^mat^ expression in pre-implantation embryos, suggesting that young X-linked genes do not undergo upregulation like old genes (Fig. 1B). In consistence to X:A readout, the X^mat^ expression of both old and young X-linked genes was significantly higher compared to the A^mat^ expression in XaXi cells of post-implantation VE and ExE (Fig. 1B). Analysis of male embryos also showed that while old X-linked genes are consistently upregulated, young X-linked genes do not (Fig. 1B). We observed that X^mat^ expression for young X-linked genes was equivalent to A^mat^ at pre-implantation, but X^mat^ expression was slightly increased compared to the A^mat^ at post-implantation (not statistically significant), indicating upregulation of these genes at post-implantation lineages.

**Figure 1.**
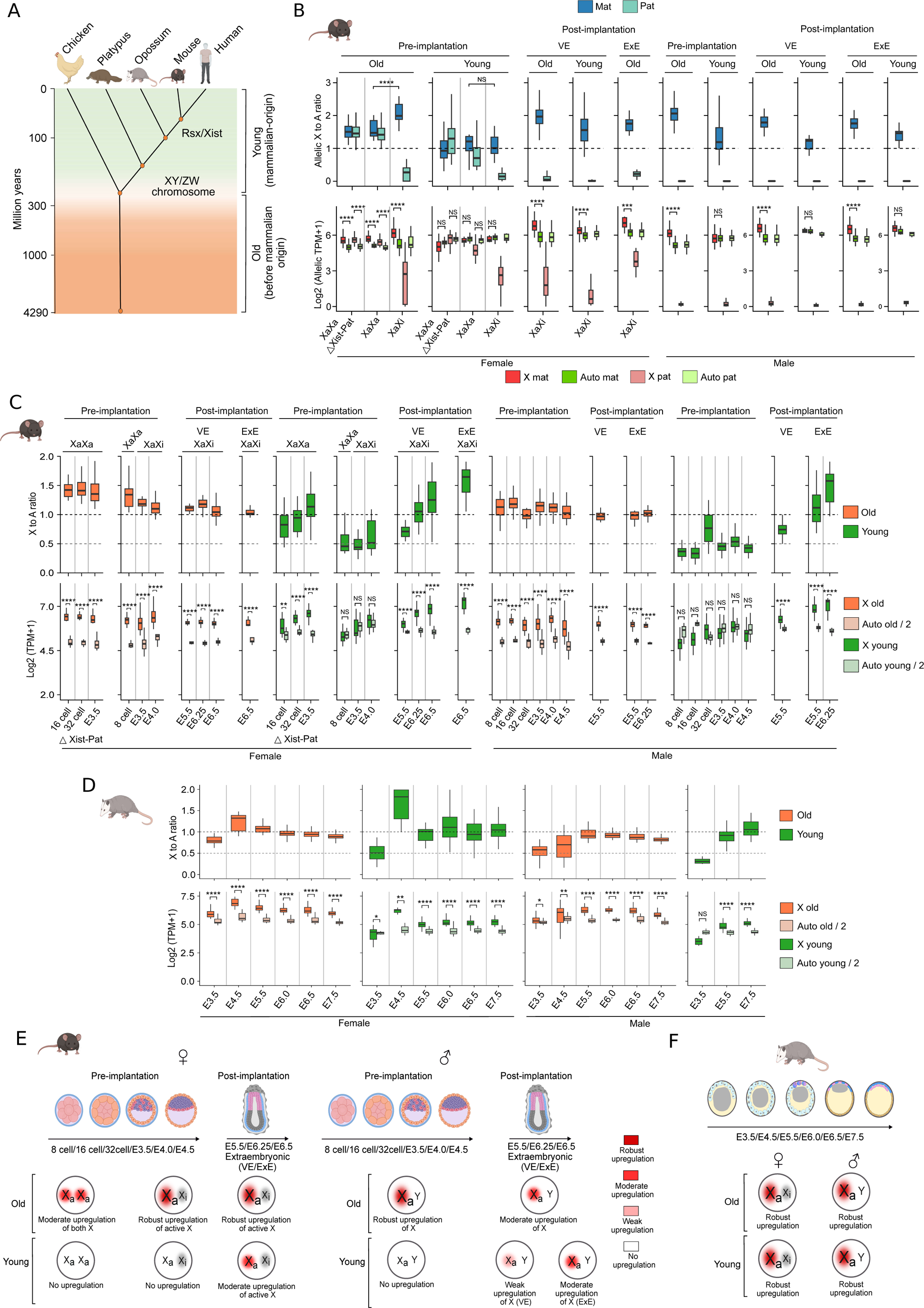
X to autosome dosage compensation dynamics of old vs. young genes during imprinted XCI in mouse and opossum embryos. (A) Schematic showing the old (genes originated before mammals) vs. young genes (mammalian origin genes). (B) Top: Plots representing the allelic X:A ratio for old vs. young genes in female (XaXa: WT/ΔXist-Pat and XaXi) and male cells of pre- and post-implantation (VE and ExE) mouse embryos. Bottom: Plots showing the allelic expression of X linked vs. autosomal genes of old and young categories in female (XaXa: WT/ΔXist-Pat and XaXi) and male cells of pre- and post-implantation (VE and ExE) mouse embryos. (C) Top: Plots depicting the non-allelic X:A ratio for old vs. young genes in female (XaXa: WT/ΔXist-Pat and XaXi) and male cells; bottom: Plots showing the expression of X linked vs. autosomal genes (Auto/2) in female (XaXa: WT/ΔXist-Pat and XaXi) and male cells of pre- and post-implantation (VE and ExE) mouse embryos. (D) Plots representing X:A ratio (top) and X-linked vs. autosomal (auto/2) expression (bottom) for old and young genes at different stages of opossum embryogenesis (male and female). (E) Model showing the active-X upregulation dynamics of old vs. young genes during the imprinted XCI at pre and post implantation (VE and ExE) mouse embryos. (F) Schematic showing the X to autosome dosage compensation pattern of old vs. young genes during opossum embryogenesis for both sexes. In each boxplots represented in the figure, the inside line denotes the median value, and the edges represent 25% and 75% of the dataset, respectively. One-sided Wilcoxon rank-sum tests: P < 0.0001; ****, < 0.001; ***, < 0.01; **, < 0.05; *, NS; non-significant.

One of the limitations of allele-specific analysis is that a subset of genes gets excluded from the analysis due to lack of qualified SNPs. Therefore, we attempted to perform XCU analysis in XaXa and XaXi cells through non-allelic approaches so that we have better representation of different classes of evolutionary genes. We excluded escapee genes for non-allelic analysis as they can express from inactive-X as well and through non-allelic approach we will not be able to disentangle their expression from active vs. inactive allele in XaXi cells. Moreover, through non-allelic approach we performed XCU analysis across different developmental stages to get better clarity on developmental dynamics of XCU. In case of X-upregulation in XaXi cells, the expected X:A ratio should be greater than 0.5 and close to 1. Indeed, we observed that while the X:A ratio for old genes in XaXi cells of pre-implantation embryos was above 1, young genes exhibited near to 0.5, further indicating robust upregulation of old genes but no upregulation of young X-linked genes at preimplantation stage (Fig. 1C). Moreover, we compared the X:A ratio of young genes of XaXi cells of E3.5 with the same stage (E3.5) XaXa (ΔXist-Pat) cells. We found that while X:A ratio of XaXa (ΔXist-Pat) cells was above 1, it remains below 0.5 in XaXi cells at E3.5, which is less than half of the X:A ratio of the XaXa (ΔXist-Pat) cells, further indicating no upregulation of young X-linked genes at E3.5 (Fig. 1C). However, in post-implantation lineages, X:A ratio of young genes becomes above 0.5 and reaches to 1 and even more, suggesting robust upregulation of young X-linked genes at post-implantation (Fig. 1C). Old genes also maintained active-X upregulation at post-implantation. Analysis of male cells exhibited similar trends (Fig. 1C). Finally, we validated our observation through comparing the X chromosome expression with the autosomal expression. For this we devided autosomal expression by 2 (denoted by A^half^) to compare the expression of one autosomal allele equivalent to one active-X present in XaXi cell or XaY cells. As expected, the X-linked expression in XaXa cells was almost doubled compared to the A^half^ at most of the data points, thereby validating our analysis (Fig. 1C). We observed that the expression of old X-linked genes in XaXi cells was always significantly higher compared to the A^half^ expression across pre- and post-implantation, further corroborating robust upregulation of old genes (Fig. 1C). However, there was no significant differences between X and A^half^ for young genes in pre-implantation embryos, reconfirming that young genes do not undergo upregulation at this stage (Fig. 1C). However, we observed that the expression of young X-linked genes was significantly higher compared to the A^half^ in post-implantation lineages, further validating active-X upregulation of young genes at post-implantation. Similar pattern was observed in male cells as well (Fig. 1C). Together, we conclude that the active-X upregulation dynamics differs between old and young X-linked genes during the imprinted XCI in mouse early embryos. While old genes undergo robust active-X upregulation starting from pre-implantation, young X-linked genes undergo active-X upregulation at post-implantation, not at pre-implantation (Fig. 1E).

### Both old and young genes undergo XCU during marsupial embryogenesis

Next, we explored the XCU dynamics of old vs. young X-linked genes in opossum embryos, which undergo imprinted form of XCI, using available scRNA-seq dataset (Mahadevaiah et al. 2020). We explored overall X:A readout (non-allelic) across early developmental stages (E3.5, E4.5, E5.5, E6.0, E6.5 and E7.5). In opossum, it has been demonstrated that XCI occurs starting from E3.5 onwards and it becomes robust at E4.5) (Mahadevaiah et al. 2020). Overall, we observed that X:A ratio for both old and young genes in female cells were above or close to ∼1 across developmental stages, indicating robust XCU upon XCI (Fig. 1D). Only at E3.5 stage, we observed that XCU was not robust like other developmental stages (Fig. 1D). Similar trend was noticed for male embryos as well (Fig. 1D). To validate further, we compared the X-linked expression with the A^half^ expression, which corroborated the similar trends as observed through the X:A ratio analysis (Fig. 1D). Taken together, we conclude that overall old and young genes undergo robust XCU during opossum embryogenesis (Fig. 1F).

### Old and young genes undergo active-X upregulation upon random XCI in post-implantation mouse embryos and differentiated ESC

Next, we investigated the active-X upregulation dynamics of old and young genes during the initiation and establishment of random XCI. For this, we performed allele-specific analysis of available scRNA-seq dataset of post-implantation (E5.5, E6.25 and E6.5) mouse embryos(Cheng et al. 2019). Allelic X:A ratio analysis as well as expression analysis revealed that both old and young genes undergo robust XCI in post-implantation embryos (Fig. 2A). Interestingly, we observed that the active-X:A ratio significantly increased in XaXp and XaXi cells compared to the XaXa cells, signifying that both classes of genes undergo active-X upregulation upon initiation of random XCI (Fig. 2A). Allelic expression analysis of X-linked and autosomal genes corroborated the same fact (Fig. 2A). Active-X upregulation was observed in male embryos as well (Fig. 2A). Furthermore, analysis of same set of genes used for the analysis of imprinted XCI in embryos also resulted similar outcomes in post-implantation epiblast cells undergoing random XCI (Fig. S2). Next, we extended our analysis to two independents female mESC differentiation datasets (Böttcher et al. 2020; Lentini et al. 2022) (Fig. 2B and 2C). ESC differentiation represents the initiation of random XCI and has been extensively used to explore mechanistic aspect of random XCI (Jonkers et al. 2009; Gontan et al. 2012; Monkhorst et al. 2008; Bowness et al. 2022; Moindrot et al. 2015). We observed significant increase of active-X:A ratio in XaXp and XaXi cells compared to the XaXa cells upon initiation of random XCI for both old and young genes, suggesting the robust active-X upregulation of both classes of genes (Fig. 2B and 2C). Allelic expression analysis also showed that active-X expression in XaXi cells for both classes of genes was always significantly higher compared to the corresponding autosomal allele, further validating their upregulation (Fig. 2B and 2C). Additionally, we observed moderate upregulation of both X in XaXa cells for old and young genes (Fig. 2B, 2C and 2D). On the other hand, in undifferentiated male ESC, while young genes showed robust upregulation, old genes showed moderate upregulation (Fig. 2B and 2D). Taken together, we conclude that both old and young genes undergo active-X upregulation upon random XCI in mouse (Fig. 2D).

**Figure 2.**
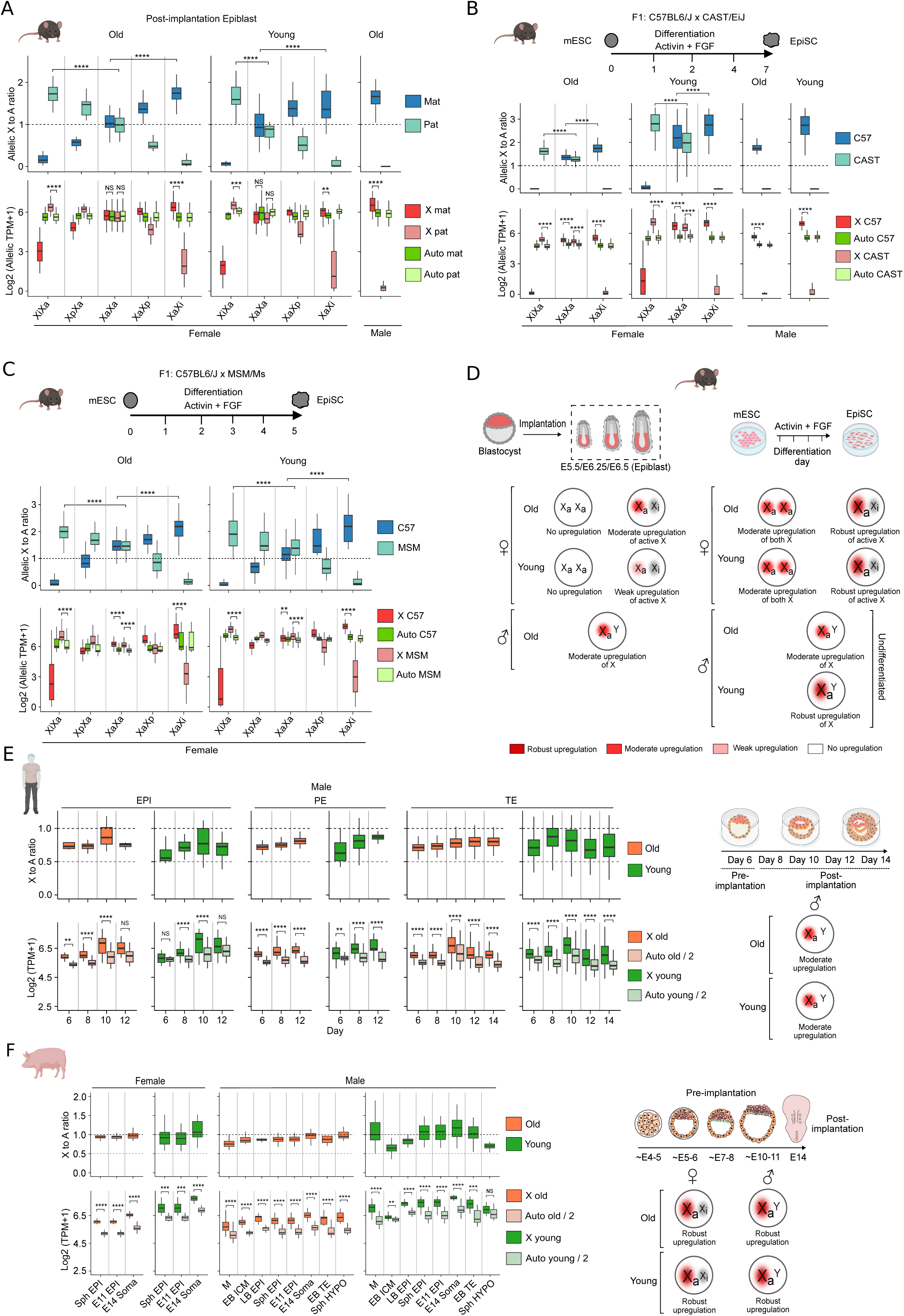
Old and young genes undergo X to A dosage compensation during random XCI in mouse as well as during human and pig embryogenesis. (A) Allelic X:A ratio (top) and allelic expression of X linked vs. autosomal genes (bottom) for old vs. young genes in female (XaXa, XaXp and XaXi) and male post-implantation epiblast of mouse. (B) and (C) Allelic X:A ratio (top) and allelic expression of X linked vs. autosomal genes (bottom) for old vs. young genes in female (XaXa, XaXp and XaXi) and male differentiated ESC (C57BL6/j X CAST/EiJ and C57BL6/j X MSM/Ms). (D) Model diagram showcasing the active-X upregulation status of XaXa, XaXi and XaY cells in mouse post-implantation epiblast and differentiated ESC. (E) X:A ratio (top) and expression of X and auto genes/2 (bottom) of old vs. young category in EPI, PE and TE lineages of male human embryos (*in vitro*) of different developmental stages. Right: Schematic presentation of X to A dosage compensation pattern during human embryogenesis (F) X:A ratio (top) and expression of X and auto genes/2 (bottom) for old vs. young genes during pig embryogenesis (male and female). Right: Model diagrams showing the X to A dosage compensation pattern in male and female pig embryos. In boxplots, the inside line: median value, and the edges represent 25% and 75% of the dataset, respectively. One-sided Wilcoxon rank-sum tests: P < 0.0001; ****, < 0.001; ***, < 0.01; **, < 0.05; *, NS; non-significant.

### Old and young genes undergo XCU during human and pig embryogenesis

Next, we explored the XCU status of old and young genes during human and pig embryogenesis. For human, we analyzed different lineages (Epiblast: EPI, Primitive endoderm: PE and Trophectoderm: TE) of *in vitro* developed male embryos of different stages (d6, d8, d10, d12 and d14) using available scRNA-seq datasets (Fig. 2E) (Zhou et al. 2019). X:A analysis revealed that both old and young genes undergo moderate upregulation of the active-X in male human embryonic lineages (Fig. 2E). Further, expression of X-linked genes (old and young) was always significantly higher compared to the A^half^ expression, signifying upregulation of the active-X in these embryos (Fig. 2E). Next, we analyzed pig embryos using available scRNA-seq datasets (Zhu et al. 2021; Ramos-Ibeas et al. 2019). For female we used spherical epiblast (Sph EPI), E11 EPI and E14 soma, which has been reported to undergo XCI (REF) (Fig. 2F). We analyzed different lineages from male embryos as well (Fig. 2F). We observed that X:A ratio of both old and young genes was close to 1 or above 1 in male as well as female, indicating robust XCU for both categories of genes (Fig. 2F). Importantly, the expression of X-linked genes (old and young) was always significantly higher compared to the A^half^ expression, reconfirming the upregulation of active-X chromosome (Fig. 2F). Taken together, we conclude that both old and young genes undergo XCU during human and pig embryogenesis.

### XCU pattern of X-orthologs and auto-orthologs differs during mouse embryogenesis

Next, we investigated the XCU dynamics based on conservation of X-linked genes between eutherian and metatherian X chromosome. For this, we classified genes into two categories: (1) X-linked genes conserved between eutherian X and metatherian X (X-orthologs) and (2) eutherian X-linked genes, which are not conserved with opossum X but orthologous to opossum autosomal genes (Auto-orthologs) (Fig. 3A). First, we investigated the active-X upregulation dynamics of these classes of genes during the imprinted XCI in mouse using allele-specific approach. Allelic X:A ratio and expression analysis revealed that both classes of genes undergo robust XCI at pre-implantation and post-implantation lineages (Fig. 3B). Simultaneously, we observed significant increase in active-X:A ratio in XaXi cells compared to the XaXa cells for both classes of genes at pre-implantation and post-implantation (VE and ExE), suggesting active-X upregulation of these genes upon XCI (Fig. 3B). Allelic expression analysis also resulted similar outcomes (Fig. 3B). Male embryos also exhibited similar trend (Fig. 3B). However, we noticed that the overall extent of active-X upregulation was differed, while X-orthologs exhibited robust XCU, XCU pattern for auto-orthologs was weak (Fig. 3B and 3E). To validate further, we performed non-allelic X:A and expression analysis across different developmental stages. For this, we removed escapee genes from our analysis. We observed that X-orthologs undergo robust active-X upregulation in XaXi cells and XaY cells across pre- and post-implantation stages (Fig. 3C). However, we observed that the extent of upregulation of auto-ortholog genes was lower and was not uniform across stages and lineages (Fig. 3C). Separately, we observed that in post-implantation VE and ExE cells exhibited almost no active-X upregulation, while allelic analysis showed upregulation (Fig. 3B and 3C). We assume that these differences could be due to the differences in set of genes between the two methods. Taken together, we conclude that the active-X upregulation pattern differs between X-orthologs and auto-orthologs during imprinted XCI in female as well as male embryos of mouse (Fig. 3E). Additionally, we observed weak upregulation of both X for both classes of genes in XaXa cells of pre-implantation embryos (Fig. 3B and 3E).

**Figure 3.**
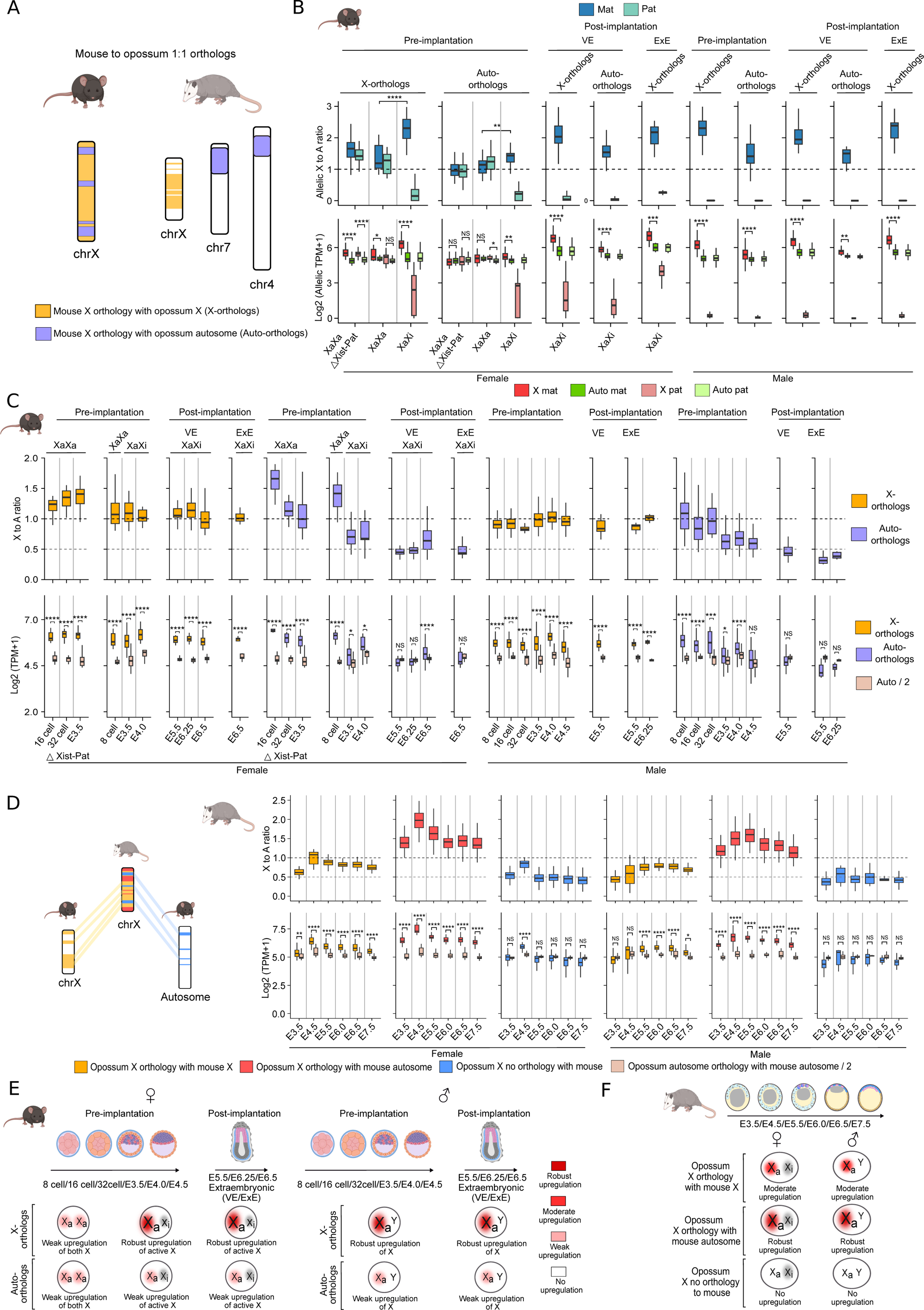
X to A dosage compensation of X-orthologs vs. auto-orthologs during imprinted XCI in mouse and opossum embryos. (A) Schematic showing the classification of X-orthologs (conserved between mouse and opossum X) and Auto-orthologs genes (mouse X-linked genes orthologus to autosomes of opossum). (B) Allelic X:A ratio plots (top) and allelic expression plots (bottom) for X-orthologs vs. auto-orthologs in female (XaXa: WT/ΔXist-Pat and XaXi) and male cells (XaY) of preimplantation and post-implantation (VE and ExE) mouse embryos. (C) X:A and expression analysis of X-orthologs vs. auto-orthologs at different stages of mouse embryogenesis (male and female). (D) Plots representing the X:A ratio (top) and expression profile (bottom) for different categories genes based on mouse and opossum ortholog at different stages of opossum embryogenesis. (E) Overview of the active-X upregulation status of X-orthologs vs. auto-orthologs in XaXa, XaXi and XaY cells during mouse pre-implantation development and post-implantation lineages (VE and ExE) undergoing imprinted XCI. (F) Schematic view of X to A dosage compensation pattern of different categories of mouse and opossum orthologs during opossum embryogenesis. One-sided Wilcoxon rank-sum tests: P < 0.0001; ****, < 0.001; ***, < 0.01; **, < 0.05; *, NS; non-significant.

Next, we extended our analysis during random XCI in mouse post-implantation epiblast and differentiated ESCs. Allelic X:A and allelic expression analysis revealed that both classes of genes undergo robust XCI during the random XCI (Fig. 4A, 4B and 4C). Simultaneously, we observed that X-orthologs undergo robust active-X upregulation upon XCI as indicated by significant increase of active-X:A ratio in XaXi cells compared to the XaXa cells in post-implantation epiblast and differentiated ESCs (Fig. 4A, 4B and 4C). Similarly, we observed that the expression of X-linked genes from the active-X was always significantly higher compared to the corresponding autosomal allelic expression in XaXi cells, further validating the upregulation of X-orthologs (Fig. 4A, 4B and 4C). Overall, male embryos depicted similar trends (Fig. 4A and 4B). On the other hand, for auto-orthologs, we observed significant increase in active-X:A ratio in XaXi cells compared to the XaXa cells in most of the cases in post-implantation epiblast and differentiated ESCs, suggesting upregulation of auto-ortho genes upon XCI (Fig. 4A, 4B and 4C). However, we observed that the overall pattern of XCU was weak. Similarly, we observed that expression of auto-ortho genes from the active-X was not significantly higher compared to the corresponding autosomal allele in many cases of XaXi cells (Fig. 4A, 4B and 4C). Together, we conclude that the pattern of active-X upregulation upon initiation of random XCI differs between X-orthologs and auto-orthologs (Fig. 4D). Separately, we observed weak upregulation of both of the X-chromosome in XaXa cells for X-orthologs genes, which was not observed for auto orthologs (Fig. 4A-D).

**Figure 4.**
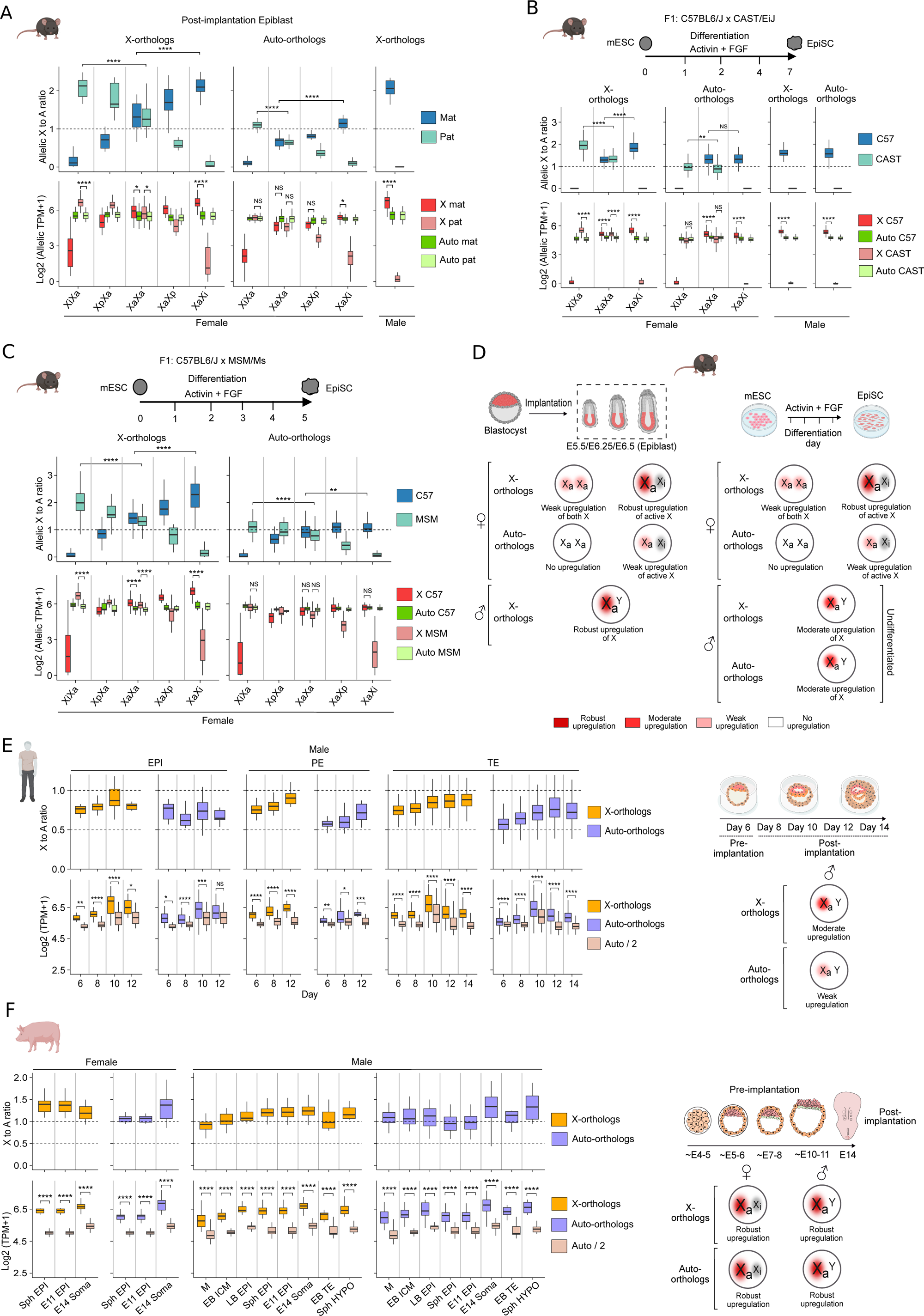
Dosage compensation pattern of X-orthologs vs. auto-orthologs during mouse, human and pig embryogenesis. (A) Top: Plots representing allelic X:A ratio for X-orthologs vs. auto-orthologs in female cells (XaXa, XaXp and XaXi) during initiation of random XCI and male cells of mouse post-implantation epiblast. Bottom: Plots representing corresponding allelic expression comparison between X and Autosomal genes of different categories. (B) and (C) presenting the allelic X:A and expression analysis of X-orthologs vs. auto-orthologs during mouse ESC differentiation. (D) Diagrammatic overview of active-X upregulation status of cells undergoing random XCI and male cells in mouse post-implantation epiblast and differentiated ESC. (E) Analysis representing the X to A dosage compensation pattern of X-orthologs vs. auto-orthologs in different lineages of human embryos (*in vitro*). (F) Plots (X:A ratio and expression of X and auto/2) demonstrating the X to A dosage compensation pattern of X-orthologs vs. auto-orthologs at the different stages of pig embryogenesis. Right: Model diagram illustrating the overall X to A balance during pig embryogenesis. One-sided Wilcoxon rank-sum tests: P < 0.0001; ****, < 0.001; ***, < 0.01; **, < 0.05; *, NS; non-significant.

### XCU pattern of X-orthologs and auto-orthologs differs across eutherian and metatherian species

Next, we explored the XCU status of X-orthologs during opossum embryogenesis. We found that these genes undergo XCU in opossum embryos, however, it was not robust as it was observed for mouse (Fig. 3D and 3F). Additionally, we observed that the extent of XCU was higher for X-linked genes of opossum orthologues to mouse autosomes (Fig. 3D and 3F). Surprisingly, we observed that X-linked genes of opossum, which do not have orthologs to either X or autosomes of mouse do not undergo XCU (Fig. 3D and 3F). On the other hand, in male human embryonic lineages we observed that XCU pattern of X-orthologs was moderate not robust like mouse (Fig. 4E). However, the pattern for auto-orthologs was weak as observed in mouse (Fig. 4E). Interestingly, in case of pig embryos, both X-orthologs and auto-orthologs exhibited robust XCU (Fig. 4F). Taken together, we conclude that the XCU pattern of X-orthologs and auto-orthologs differs across eutherian and metatherian species.

### Evolutionary stratum-specific differences in X to A dosage compensation

Next, we investigated the dosage compensation pattern of X-linked genes based on their distinct evolutionary stratum as reported previously (Lahn and Page 1999). In brief, the X and Y chromosome in mammals evolved from a homologous pair of autosomes. During the evolution, there was loss of recombination between X and Y, and Y chromosome underwent multiple rounds of inversion and deletion process (Graves 2016). Based on the X and Y divergence pattern, the X chromosome in human has been demarcated into four distinct evolutionary strata (Fig. 5A). To understand the dosage compensation dynamics of X-linked genes in relation to these evolutionary strata, we identified mouse X-linked genes orthologous to human X chromosome and profiled in which strata of human X these genes belong to. Based on this, we classified genes into three categories: strata1, strata2 and strata 3+4 (genes located on strata 3 and strata 4) (Fig. 5A). Allelic X:A ratio analysis as well as allelic expression analysis revealed that all these three categories of genes undergo robust imprinted XCI (Fig. 5B). On the other hand, we observed that the allelic X:A ratio (for both paternal and maternal-X) of strata1 and strata 2 genes was above 1 in XaXa cells (WT and ΔXist-Pat) of pre-implantation embryos, indicating hyper activation of both of this X-chromosome (Fig. 5B). Allelic expression analysis also showed significant higher expression of both paternal and maternal-X compared to the autosomal allele in XaXa cells (Fig. 5B). Interestingly, allelic X:A ratio of maternal-X was significantly higher than the paternal-X in XaXa (WT) cells for strata1 genes, suggesting upregulation of these genes possibly precedes XCI (Fig. 5B). In XaXi cells, the allelic X:A ratio for maternal-X maintained above 1 for strata1 genes indicating upregulation of active-X (Fig. 5B). However, we did not see any further significant increase in X:A ratio in XaXi cells compared to the XaXa cells, further validating that maternal X upregulation was already in place in XaXa cells even before robust inactivation of X-linked genes for strata1 genes (Fig. 5B). However, for strata2 genes, we observed significant increase in maternal active-X:A in XaXi cells compared to the XaXa cells, suggesting active-X upregulation upon XCI (Fig. 5B). Additionally, upregulation of active-X for strata1 and strata2 genes was maintained in XaXi cells of post implantation VE and ExE cells (Fig. 5B). In males, we observed active-X upregulation of strata1 and strata2 genes in pre-implantation and post-implantation lineages, except for strata2 genes in ExE cells (Fig. 5B). Next, we validated our observation across different stages of development through non-allelic approach, where we excluded potential escapee genes from our analysis as their expression also originate from inactive allele. In consistent with the allelic analysis, we observed that strata1 genes undergo robust X to A dosage compensation in pre- and post-implantation embryos of both sexes (Fig. 5C). But for strata2 genes we observed that in pre-implantation embryo X to A dosage compensation was not robust like strata 1 in female (XaXi) as well as in male. In post-implantation, we observed robust X to A dosage compensation in female but not in male (Fig. 5C). On the other hand, we found that the active-X upregulation pattern of strata3+4 genes differed from strata1 and strata2 genes. We observed that strata 3+4 genes undergo active-X upregulation in post-implantation embryo (VE) but not in pre-implantation of female cells (Fig. 5B). In contrary, male pre-implantation embryos showed weak active-X upregulation (Fig. 5B). Through non allelic approach with exclusion of escapee showed that strata 3+4 genes do not undergo X to A dosage compensation in pre and post implantation embryos of both sexes (Fig. 5C). Together, we conclude that XCU pattern in mouse differs between X-linked genes belonging to different strata of human X during imprinted XCI (Fig. 5D).

**Figure 5.**
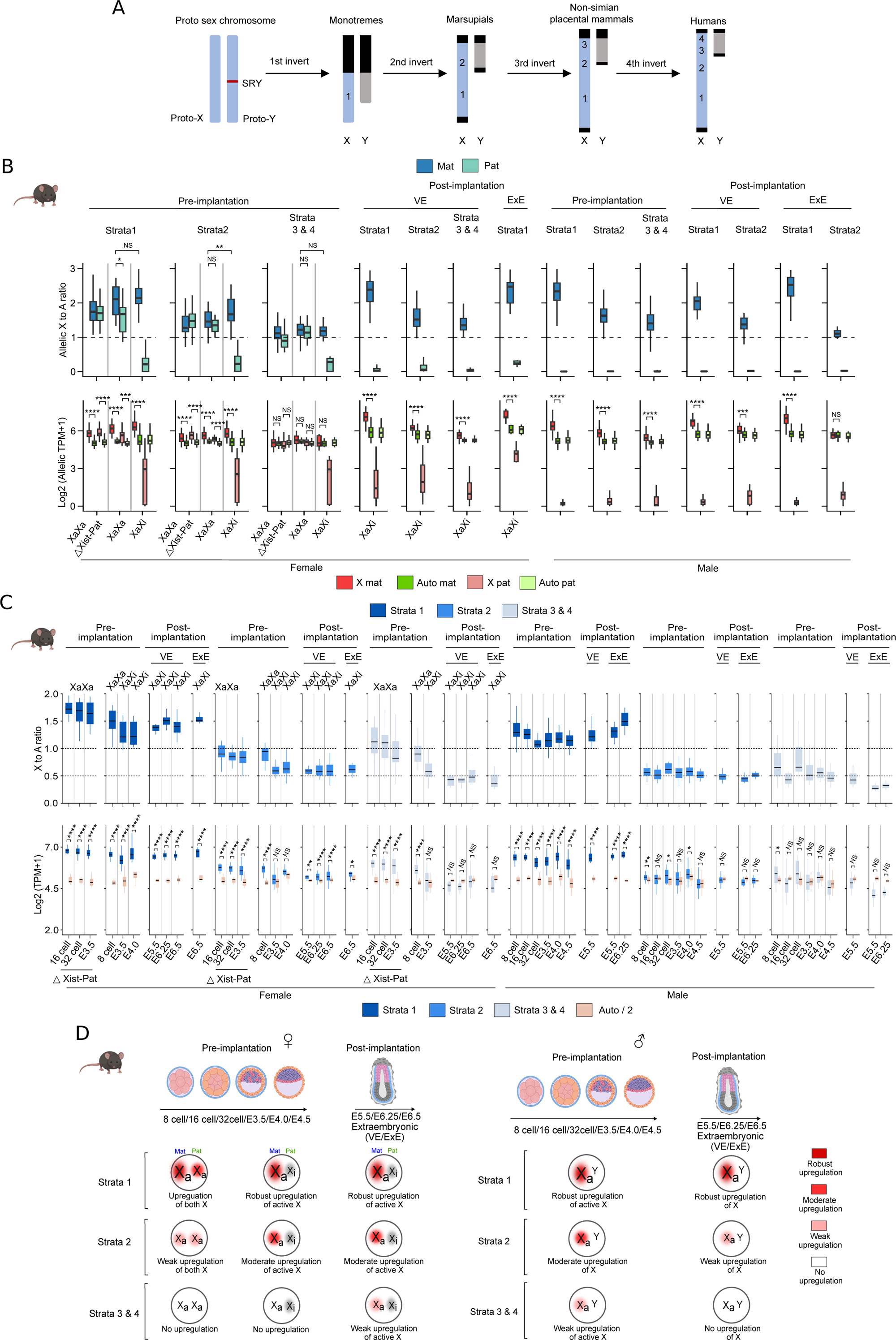
Active-X upregulation dynamics of different evolutionary strata genes during imprinted XCI in mouse embryos. (A) Illustration of different evolutionary strata (Strata 1, 2, 3 and 4) of human X-chromosome. (B) Plots showing the active-X upregulation analysis of X-linked genes of different strata in mouse pre- and post- implantation cells with imprinted XCI: allelic X:A analysis (top) and allelic expression analysis (bottom). (C) represents similar analysis but without allelic resolution across different stages of pre and post implantation development. (D) Schematic showing the active-X upregulation dynamics of genes belonging to different strata during imprinted XCI in mouse embryos. One-sided Wilcoxon rank-sum tests: P < 0.0001; ****, < 0.001; ***, < 0.01; **, < 0.05; *, NS; non-significant.

Next, we investigated what happens during the random XCI. We noticed that all three classes of genes (strata1, strata2 and strata 3+4) undergo robust XCI during the random XCI in post-implantation mouse epiblast and differentiated ESC (Fig. 6A, 6B and 6C). We observed that active X:A ratio of strata1 and strata2 genes was significantly increased in XaXi cells compared to the XaXa cells in post-implantation epiblast and differentiated ESC, suggesting they undergo active-X upregulation upon random XCI (Fig. 6A, 6B and 6C). Allelic expression analysis corroborated the same (Fig. 6A, 6B and 6C). However, we observed that while strata 1 genes exhibit robust XCU, XCU pattern of strata 2 genes were moderate or weak (Fig. 6D). On the other hand, in most of the cases, active-X:A ratio of strata 3 +4 genes increased in XaXi cells compared to the XaXa cells of epiblast and differentiated ESC, indicating upregulation of these genes upon XCI. However, in most of the cases the expression of active-X genes was not significantly higher compared to the corresponding autosomal allele and even some cases it was lower than the autosome. Together, we conclude that strata3+4 genes either do not undergo upregulation or upregulation is very weak during the random XCI in mouse (Fig. 6D).

**Figure 6.**
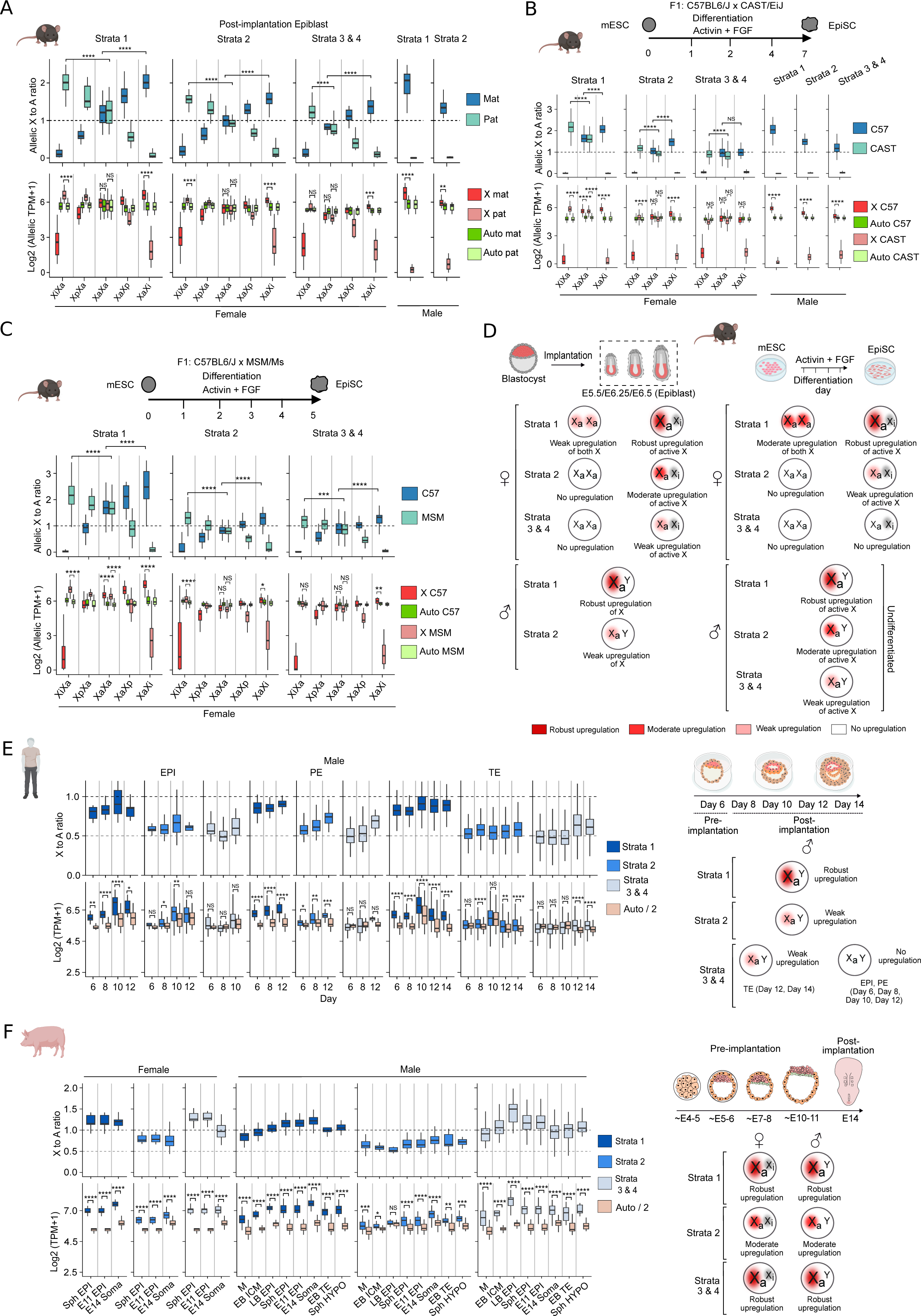
X to A dosage compensation dynamics of different evolutionary strata genes during random XCI in mouse and during human and pig embryogenesis. (A) Strata-wise active-X upregulation dynamics during random XCI in female mouse post-implantation epiblast cells as well as in male epiblast cells; top: allelic X:A analysis and bottom: allelic expression analysis. (B) and (C) represents strata-wise X upregulation dynamics during mouse ESC differentiation. (D) Schematic depiction of active-X upregulation pattern during initiation of random XCI in female mouse post-implantation epiblast and differentiated ESC alongside in male cells (XaY). (E) X to A dosage compensation pattern of X-linked genes of different strata in male human embryos. Top: X:A ratio quantification, bottom: Expression analysis of X linked genes of different strata vs. autosomal genes. Right: Schematic overview of dosage compensation pattern. (F) X to A dosage compensation profiling of different evolutionary strata genes during pig embryogenesis through X:A ratio (top) and expression analysis (bottom). Right panel representing a schematic overview of X to A dosage compensation pattern during pig embryogenesis. One-sided Wilcoxon rank-sum tests: P < 0.0001; ****, < 0.001; ***, < 0.01; **, < 0.05; *, NS; non-significant.

Next, we extended our analysis for male human embryos (EPI, PE and TE). X:A ratio for strata 1 genes revealed moderate to robust upregulation (Fig. 6E). Further, expression of X-linked genes was always significantly higher compared to the haploid set of autosomes (Auto^half^), confirming the upregulation of strata1 genes (Fig. 6E). However, the X:A ratio for strata 2 genes was above 0.5 but lesser than 0.75 in most cases indicating weak upregulation of this class of genes (Fig. 6E). Expression analysis revealed that expression of X was not always significantly higher compared to the Auto^half^ further corroborating weak upregulation (Fig. 6E). Interestingly, we observed that strata3+4 genes do not undergo upregulation except in d12 and d14 TE cells (Fig. 6E). Together, we conclude that dosage compensation pattern of genes of different evolutionary strata differs during human embryogenesis like mouse. In contrary, we observed that all three strata classes undergo XCU during pig embryogenesis. Specially, X:A ratio for strata1 and strata 3+4 genes was close to 1 or above 1 indicating robust upregulation of these genes (Fig. 6F). However, strata2 genes exhibited weak to moderate upregulation (X:A ∼ 0.5-0.75) (Fig. 6F).

### Enrichment of active-chromatin marks correlates with XCU pattern of different evolutionary classes of genes

Next, we explored the underlying molecular basis of differential XCU pattern of different evolutionary classes of genes. Recently, chromatin accessibility and enrichment of active marks have been implicated to be associated with XCU (Allsop et al. 2025; Talon et al. 2021; Naik et al. 2022). Therefore, we asked whether enrichment of active chromatin marks is linked to the differential XCU pattern observed for different evolutionary classes of genes. To address this, we profiled the enrichment of H3K4me3, H3K4me1 and H3K27ac around transcriptional start sites (TSSs) of X-linked genes relative to autosome (Chr-13) using available target chromatin indexing and tagmentation (TACIT) datasets of mouse preimplantation embryo (blastocyst) (Liu et al. 2025). Similarly, we measured the enrichment for H3K36me3 across gene body, which involves in transcriptional elongation (Liu et al. 2025). For autosomal genes, we divided the reads by 2 to make it equivalent to one active-X chromosome of male and female. Interestingly, we observed that old X-linked genes, which undergo robust upregulation were enriched with H3K4me1, H3K27ac and H3K36me3 relative to the old Chr13 genes in blastocyst (Fig. 7A and Fig. 8). However, there was no significant differences observed for enrichment of H3K4me3 between X and Chr13 for old genes (Fig.7A). On the other hand, young genes which do not undergo XCU at pre-implantation, did not exhibit such enrichments of these marks relative to young Chr13 genes (Fig. 7A and Fig. 8). Together, we conclude that the differential X-upregulation pattern of old vs. young X-linked genes at pre-implantation possibly driven by the chromatin state. Similarly, we observed that X-ortholog, strata1 and strata2 genes, which undergo moderate to robust XCU were enriched with H3K4me1, H3K27ac and H3K36me3 relative to the old Chr13 genes, but not for H3K4me3 (Fig. 7B, 7C and Fig. 8). However, auto-ortho and strata3+4 genes, which exhibit weak or no XCU, showed higher enrichment of H3K4me1 and H3K27ac compared to the Chr13, but not for H3K4me3 and H3K36me3 (Fig. 7B, 7C and Fig. 8). Taken together, we conclude that differential pattern of active-X upregulation of different evolutionary classes of genes are linked to differential enrichment of different active marks in pre-implantation embryo.

**Figure 7.**
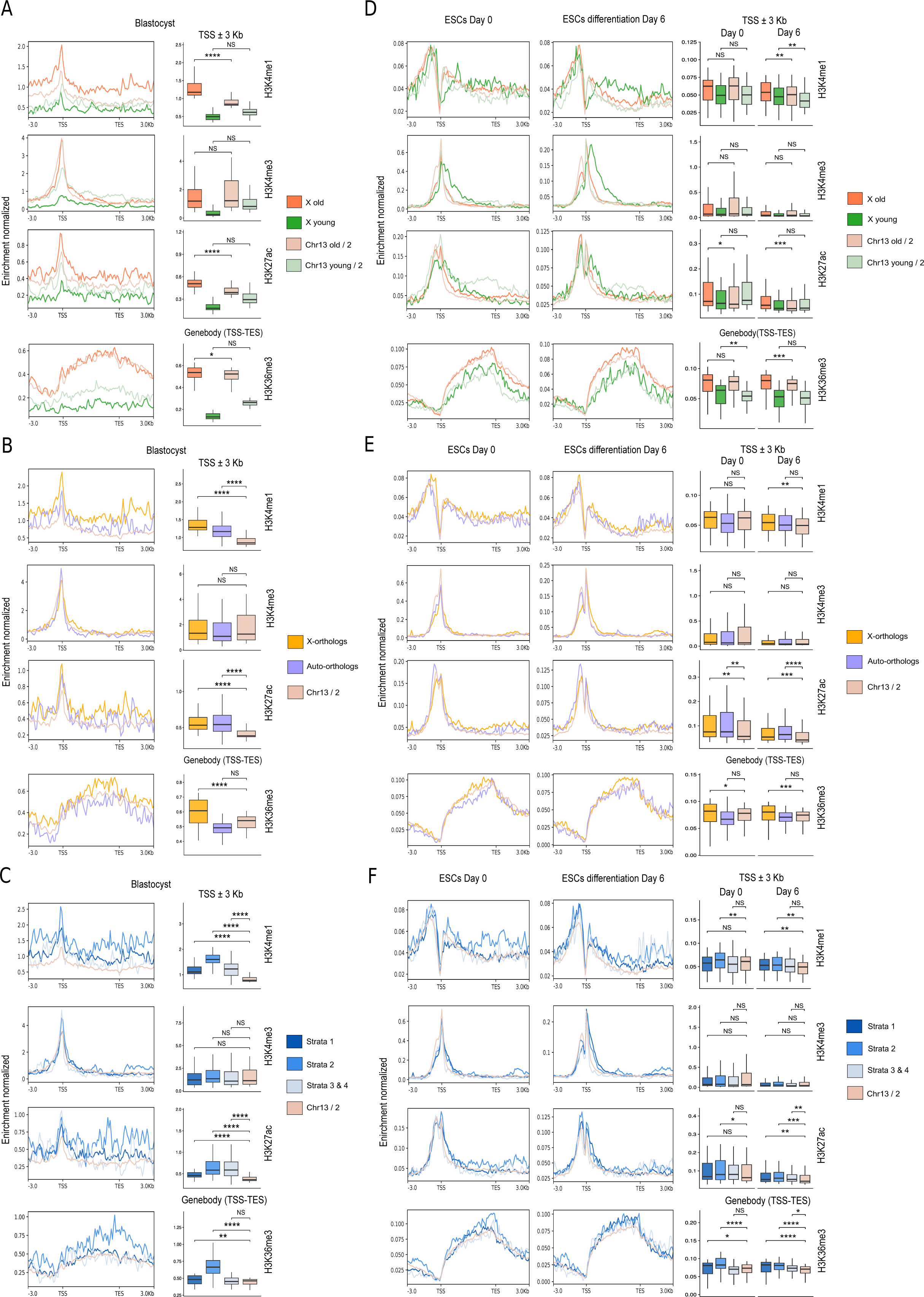
Active chromatin marks enrichment often linked to the XCU pattern. Quantification of enrichment of different active marks (H3K4me1, H3K4me3, H3K27ac and H3K36me3) on different evolutionary classes of X-linked genes relative to autosomal (Chr13) genes in blastocyst: (A) old and young genes (B) X-orthologs and auto-orthologs (C) Strata1, strtata2 and strata 3+4. (D) (E) and (F) plots showing enrichment of different active marks (H3K4me1, H3K4me3, H3K27ac and H3K36me3) on different evolutionary classes of X-linked genes relative to autosomal (Chr13) genes in d0 ESC and d6 differentiated ESC. One-sided Wilcoxon rank-sum tests: P < 0.0001; ****, < 0.001; ***, < 0.01; **, < 0.05; *, NS; non-significant.

**Figure 8.**
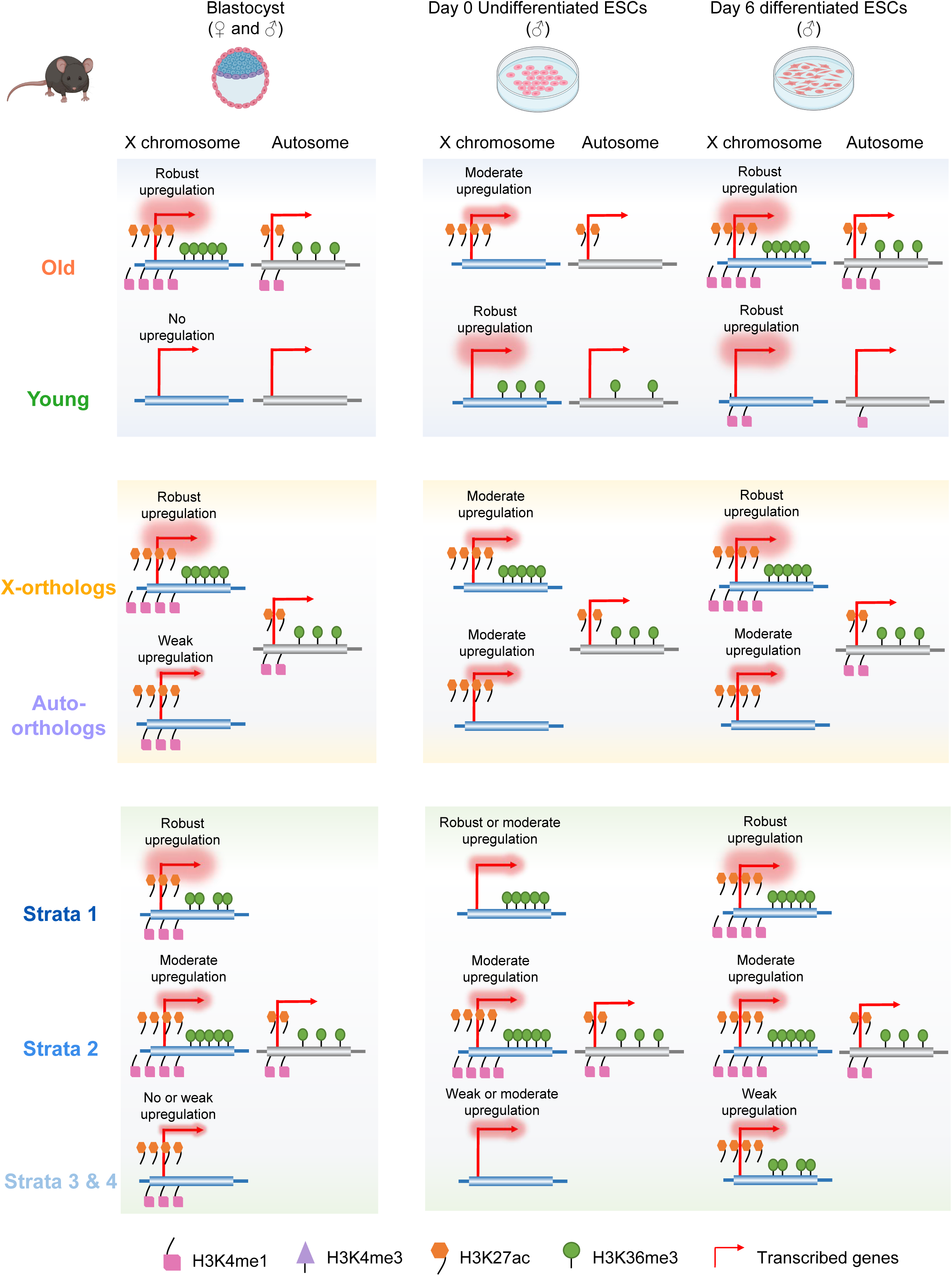

Next, we extended our analysis to undifferentiated and differentiated male ESC using available ChIP-seq datasets (Xiao et al. 2012). In this undifferentiated male ESC, while old genes exhibit moderate XCU, young genes undergo robust XCU and they exhibited differential enrichment pattern of active marks (Fig. S3A). While old genes were enriched for H3K27ac, young genes showed enrichment of H3K36me3 relative to autosome (Fig. 7D). Separately, X-ortholog and auto-ortho genes, which undergo moderate XCU, showed enrichment for H3K27ac (Fig. 7E, Fig. S3B). However, X-orthologs showed enrichment for H3K36me3 as well. Among the different strata classes, strata2 had greater enrichment for H3K4me1, H3K27ac and H3K36me3. However, strata1 showed enrichment of only H3K36me3 (Fig. 7E). On the other hand, in differentiated ESC (d6), we observed that old genes, X-ortholog and strata 1+2 genes with moderate to robust XCU pattern were significantly enriched with H3K4me1, H3K27ac and H3K36me3 relative to Chr-13 (Fig. 7D, 7E, 7F and Fig. S3). However, young genes, which undergo robust XCU showed greater enrichment for H3K4me1, (Fig. 7D and Fig. S3A). Moreover, auto-ortho, strata3+4 genes which exhibit weak to moderate XCU showed greater enrichment for H3K27ac (Fig. 7E, 7F and Fig. S3B and S3C). Additionally, strata3+4 genes were enriched with H3K36me3 as well. Together, we conclude that often the XCU can be driven by enrichment of different set of active-marks based on the evolutionary and developmental context.

## Discussion

In this study, we have explored the dosage compensation dynamics of genes with different evolutionary origins during eutherians (Mouse, human and pig) and marsupial (opossum) embryogenesis. Overall, we demonstrate that the dosage compensation pattern varies with the evolutionary origins as well as across species within a given evolutionary stratified gene group. Our study provides significant insight into the evolutionary connection of dosage compensation in eutherians and marsupials.

We show that the pattern of active-X upregulation differs between old and young genes during the imprinted XCI in early mouse embryos. Old genes are upregulated from the very beginning of imprinted XCI at preimplantation female embryos in mouse, whereas young genes do not undergo active-X upregulation at pre-implantation rather it is upregulated in post-implantation lineages (VE and ExE) (Fig. 1E). Even, male embryos follow the similar route suggesting that the dynamics of X to A dosage compensation is linked to the evolutionary age in the context of imprinted XCI in mouse (Fig. 1E). However, both old and young genes undergo XCU during imprinted XCI in female embryos of opossum (Fig. 1F). Similar trend was observed for male embryos of opossum (Fig. 1F). On the other hand, we demonstrate that both old and young genes undergo XCU during random XCI in mouse post-implantation epiblast and differentiated ESC (Fig. 2D). Moreover, we observed stable X to A dosage compensation of old and young genes during human and pig embryogenesis as well (Fig. 2E and 2F).

On the other hand, we observed that conserved genes between mouse X and opossum X (X-orthologs) undergo robust active-X upregulation in female (imprinted and random XCI) and male throughout pre-and post-implantation development in mouse (Fig. 3E and 4D). Similar trend was observed in differentiated ESC (Fig. 4D). We found similar pattern in case of opossum, human and pig for the X-orthologs; however, extent of upregulation differed across species (Fig. 3F, 4E and 4F). Specially, the XCU pattern of X-orthologs in opossum and human was not as robust as it was observed in mouse and pig. On the other hand, we found that the XCU pattern of genes of eutherian-X, which are not conserved in metatherian X but are present in autosome of metatherian (auto-orthologs) differs from the X-orthologs. Specifically, auto-orthologs exhibit weak XCU in mouse (Fig. 3E and 4D). Similarly, dosage compensation pattern differs between X-orthologs and auto-orthologs in case of human (Fig. 4E). However, we observed that both classes of genes undergo robust XCU during pig embryogenesis (Fig. 4F). Together, we conclude that the dosage compensation pattern of X-orthologs and auto-orthologs varies across eutherian and metatherian species during embryogenesis.

Next, we demonstrate that genes belonging to different evolutionary strata exhibit differential XCU pattern during mouse, human and pig embryogenesis. We find that strata 1 genes always undergo robust XCU in mouse, human and pig (Fig. 5D and 6D, 6E and 6F). Whereas strata2 genes exhibited weak to moderate XCU during mouse, human and pig embryogenesis (Fig 5 and 6). On the other hand, strata 3+4 often exhibit no upregulation or weak upregulation in mouse and human except pig where it undergoes robust upregulation, suggesting differential pattern of XCU among species (Fig. 5 and 6).

Next, we demonstrate that the enrichment of active-chromatin marks are often linked to the XCU in mouse. We show that old, X-ortholog and strata1+2 genes, which undergo moderate to robust XCU in mouse blastocyst and differentiated ESC are always enriched with active chromatin marks like H3K4me1, H3K27ac and H3K36me3 relative to autosome (Fig. 8). Together, we conclude that enhanced enrichment of active-marks in these classes of genes possibly mediates upregulation. In fact, a previous study indicated that in mouse embryonic fibroblast cells (MEF) active-X are enriched with H3K4me3, RNA-polII and H3K36me3 relative to autosome (Yildirim et al. 2012). However, we did not observe enrichment of H3K4me3 in any of the evolutionary classes of genes. Furthermore, it was shown that often BRD4 and H3K27ac are enriched on the loci of upregulated X-linked genes and they play an important role in mediating XCU (Lyu et al. 2022). Moreover, a recent study depicted that there was increased accumulation of H3K9ac on the X undergoing upregulation (Allsop et al. 2025). In contrary, a recent study did not find differences of active-marks enrichment on the X between prior and after XCU (Lentini et al. 2022). Nevertheless, in drosophila, it is well established that enrichment of H4K16ac leads to the upregulation of X-chromosome in male, further signifying the role of chromatin in mediating X upregulation (Gelbart et al. 2009). On the other hand, we find that often the enrichment of active marks corelates with the differential XCU pattern of different evolutionary classes of genes in mouse. Specially, we find that young genes, which do not undergo XCU at pre-implantation blastocyst did not exhibit enrichments of any of these active marks relative to the autosome (Fig. 7A and Fig. 8). Furthermore, auto-ortho or strata 3+4 genes, which often do not undergo robust XCU at pre-implantation, were enriched with H3K4me1, H3K27ac but not with H3K36me3 in blastocyst, indicating H3K36me3 enrichment corelates with the extent of XCU (Fig. 8). Separately, in differentiated male ESC we observed that young, auto-ortho, strata3+4 were not enriched with all the three active marks (H3K4me1, H3K27ac and H3K36me3) although they undergo frequent upregulation (Fig. 8). Albeit, young genes were enriched with H3K4me1, whereas auto-ortho were only enriched with H3K27ac and strata3+4 were enriched with H3K27ac and H3K36me3 (Fig. 8). Moreover, the enrichment pattern often differed between differentiated vs. undifferentiated ESC (Fig. 8). Together, we conclude that XCU mediated by enrichment of active-marks, however, enrichment of set of active marks differs with evolutionarily class as well as with developmental context.

## Materials and Method

### Data Availability

We used publicly available scRNA-seq, bulkRNA-seq and ChIP-seq datasets for our study. Mouse pre-implantation scRNA-Seq data was obtained from GEO: GSE45719 (Deng et al. 2014), GEO: GSE80810 (Borensztein et al. 2017b), GEO: GSE89900 (Borensztein et al. 2017a), and GEO: GSE74155 (Chen et al. 2016). Mouse post-implantation scRNA-Seq data was retrieved from GEO: GSE109071 (Cheng et al. 2019). scRNA-Seq datasets of ESC differentiation was obtained from ArrayExpress: E-MTAB-9324 (Lentini et al. 2022) and DDBJ: DRA010828 (Böttcher et al. 2020). scRNA-seq dataset of human embryos was retrieved from GEO: GSE109555 (Zhou et al. 2019). scRNA-seq datasets for different stages of pig embryogenesis was obtained from GEO: GSE155136 and GSE112380 (Zhu et al. 2021; Ramos-Ibeas et al. 2019). scRNA-seq data for opossum embryonic development was retrieved from ArrayExpress: E-MTAB-7515 (Mahadevaiah et al. 2020). Target chromatin indexing and tagmentation (TACIT) datasets for blastocyst was obtained from GEO: GSE235109 (Liu et al. 2025). ChIP-seq and bulkRNA-seq datasets for E14 ESC differentiation (d0 and d6) was gathered from GEO: GSE36114 (Xiao et al. 2012). Few datasets, where the sex information of the embryos or cells were not readily available, sex was determined based on Y-linked gene counts as per the method described in corresponding literatures.

### Gene classifications based on evolutionary origins

Old (before mammalian origin) and young genes (mammalian-origin) for each species (mouse, human, pig and opossum) were classified based on the gene age date information available from the GenOrigin database (http://genorigin.chenzxlab.cn/) (Tong et al. 2021). X-orthologs (conserved) and auto-ortholog (non-conserved) genes were classified based on orthology analysis (one to one) using the Ensembl BioMart tool. Eutherian X-linked genes orthologus to marsupial X-chromosome was classified as X-orthologs. On the other hand, eutherian X-linked genes orthologus to marsupial autosomes (mostly belonging to autosome 4 and 7) was classified as auto-orthologs. Other orthologs based classification (between marsupial and eutherian) was carried out following the same method. Strata-wise classification of genes (Strata1, strata2. Strata3+4) were performed based on the defined evolutionary strata of human X chromosome (Lahn and Page 1999; Pandey et al. 2013). First, we obtained the human X-orthologs of mouse X-linked genes and then identified in which strata of the human X those genes belong to.

### RNA-seq analysis

For scRNA-seq and bulkRNA-seq analysis, reads were mapped to corresponding reference genome of each species (mouse: mm10, opossum: ASM229v1, and pig: release-113) available in Ensembl. Mapping was performed using RSEM tool (Li and Dewey 2011). Gene expression was deduced using normalized transcripts per million (TPM) method.

### Allele-specific gene expression analysis

We performed allele-specific analysis of the scRNA-seq data of mouse embryos and ESC by leveraging SNPs present in their genome as these embryos/cells were derived from the cross of divergent mouse strains. For allele-specific analysis we followed the method as described previously (Majumdar et al. 2025; Ayyamperumal et al. 2024; Naik et al. 2021, 2024, 2022). Briefly, we constructed strain-wise insilico reference genomes by inserting the strain specific SNPs into the mouse reference genome (mm10). Strain-wise SNPs was generated from the VCF file for each strains (CAST/EiJ, MSM/Ms) available at mouse genome project (https://www.sanger.ac.uk/science/data/mouse-genomes-project). Next, RNA-seq reads were mapped to the respective strain-specific insilico reference genome using STAR aligner with parameter --multimapNMax 1 (Dobin et al. 2013). Next, to obtain allelic expression for individual genes we applied following filters to select qualified SNPs and genes. We considered those SNPs as qualified SNPs, whose total read counts (Allele A + Allele B) was at least 3 to avoid any false positive. Further, those genes were considered for the allelic analysis, which harbored at least two such qualified SNPs. Allelic reads for each gene were calculated by taking average of SNP- wise reads. Allelic expression ratio was calculated for individual genes using the following formula: (Allele A or B/Allele A + Allele B). Next, to calculate allele-specific expression, we multiplied the non-allelic TPM value with the allelic ratio.

### Profiling of X to Autosome ratio

X to autosome ratio (X:A) was determined at allelic level as well as non-allelic level. Overall, we followed the method as described previously (Naik et al. 2025, 2024). First, to mitigate any bias in X:A ratio analysis due to the presence of a subsets of genes which are expressed at very low level, we selected those genes for our analysis, which had expression > 1 TPM in individual cell. Similarly, we eliminated genes with very high expression like those were above 98^th^ percentile in each individual cell. Additionally, we selected those cells for our analysis which exhibited expression of at least 10 genes, which were qualified through the above-mentioned threshold. Additionally, one of the limitations in such X:A analysis appears to be that often X:A ratio can become skewed and may not reflect true measurement because of the low numbers of X-linked genes vs. the large numbers autosomal genes. To mitigate this, we employed a bootstrapping method as described previously (Naik et al. 2024, 2025). Basically, to calculate the X:A ratio, we randomly selected a subset of autosomal genes in equal numbers to X-linked genes and we bootstrapped for 1000 times. Finally, the median value from the 1000 bootstrapped ratio was calculated to deduce the X:A ratio. For calculating the X:A ratio using bulkRNA-seq data, similar approach was taken and genes with <1 TPM and above 90^th^ percentile thresholds were excluded for the analysis. On the other hand, in case of X:A ratio calculation without allelic resolution, X-inactivation escapee genes were removed from the analysis, since these genes express from both active-X as well as the inactive-X. For mouse, a consolidated list of escapees was made from the previous reports (Borensztein et al. 2017b; Lentini et al. 2022; Reinius et al. 2010, 2016; Soh et al. 2014; Yang et al. 2010). Additionally, for human, we removed X and Y gametologs as well as X-linked genes located to psedoautosomal regions (PAR) (Lahn and Page 1999; Sandstedt and Tucker 2004). Escapee genes for pig and opossum were obtained from the previous studies (Balaton et al. 2021; Rodríguez-Delgado et al. 2014; Zou et al. 2019).

### ChIP-seq analysis

ChIP-seq analysis was performed following the method as described earlier (Majumdar et al. 2025; Ayyamperumal et al. 2024). Breifly, Fastq reads were mapped to the GRCm38 (mm10) reference genome using the Bowtie2 (Langmead and Salzberg 2012). Using BEDTools (v2.30.0), blacklisted regions were excluded from our analysis as described in the ENCODE consortium (Quinlan and Hall 2010). Data was normalized following the BPM normalization method using DeepTools *bamCoverage (Quinlan and Hall 2010)*. Enrichment analysis was performed using DeepTools function DeepTools *computeMatrix* (scale-regions -bs 100 --missingDataAsZero -m 5000 --upstream 3000 --downstream 3000) and plots were generated using DeepTools *plotProfile*. For the TACIT datasets of blastocyst enrichment analysis was performed similar way using the available *big-wig* files.

### Statistical analysis

All statistical analysis presented in this study were performed using R. For sll analysis we performed a one-sided Wilcoxon rank-sum test. Data related to allelic X:A analysis, we compared whether the X:A ratio of active-X allele in XaXi cells was greater than the XaXa cells. In Fig. 6B data, we compared whether X:A ratio of maternal allele was greater the paternal allele. For comparison of X-linked vs. autosomal gene expression, we measured whether X-linked gene expression was statistically higher compared to the autosome. Similarly, for enrichment analysis of different histone marks, we profiled whether enrichment on X-chromosome was higher compared to the autosome. All plots were made using the ggplot2 library.

## Acknowledgement

We would like to acknowledge the support from the high-performance computing (HPC) cluster of “Beagle” in Indian Institute of Science (IISc). This study is supported by the Department of Biotechnology (DBT), Govt. of India grant (BT/PR49796/MED/97/644/2023), Anusandhan National Research Foundation (ANRF), Government of India (CRG/2022/008605). We acknowledge the Bio render for preparation of model figures.

## Author contributions

SG supervised and acquired funding for this study. SG and HCN conceptualized the study. HCN performed all the bioinformatic analysis and data interpretation. PN contributed supervisory guidance. HCN and SG wrote the manuscript. All authors approved the final version of the manuscript.

## References

Al Nadaf S, Deakin JE, Gilbert C, Robinson TJ, Graves JAM, Waters PD. 2012. A cross-species comparison of escape from X inactivation in Eutheria: Implications for evolution of X chromosome inactivation. Chromosoma 121: 71–78.

Alfeghaly C, Rougeulle C. 2025. X chromosome inactivation in mammals: general principles and species-specific considerations. EMBO Rep 26: 3478–3490.

Allsop RN, Boeren J, Tan BF, Merzouk S, Poovanthingal S, van IJcken WFJ, Demmers JAA, Mira Bontenbal H, Gontan C, Gribnau J, et al. 2025. X-chromosome upregulation operates on a gene-by-gene basis at RNA and protein levels. Nature Communications 2025 16:1 16: 8352-. https://www.nature.com/articles/s41467-025-64195-3 (Accessed December 15, 2025).

Ayyamperumal P, Naik HC, Naskar AJ, Bammidi LS, Gayen S. 2024. Epigenomic states contribute to coordinated allelic transcriptional bursting in iPSC reprogramming. Life Sci Alliance 7. https://www.life-science-alliance.org/content/7/4/e202302337 (Accessed April 18, 2025).

Balaton BP, Fornes O, Wasserman WW, Brown CJ. 2021. Cross-species examination of X-chromosome inactivation highlights domains of escape from silencing. Epigenetics Chromatin 14.

Borensztein M, Saitou M, Syx L, Diabangouaya P, Okamoto I, Chen C-J, Surani A, Heard E, Picard C, Servant N, et al. 2017a. Contribution of epigenetic landscapes and transcription factors to X-chromosome reactivation in the inner cell mass. Nat Commun 8.

Borensztein M, Syx L, Ancelin K, Diabangouaya P, Picard C, Liu T, Liang J Bin, Vassilev I, Galupa R, Servant N, et al. 2017b. Xist-dependent imprinted X inactivation and the early developmental consequences of its failure. Nat Struct Mol Biol 24: 226–233. 10.1038/nsmb.3365.

Böttcher M, Tada Y, Moody J, Kondo M, Ura H, Abugessaisa I, Kasukawa T, Hon C-C, Nagao K, Carninci P, et al. 2020. Single-cell transcriptomics, scRNA-Seq and C1 CAGE discovered distinct phases of pluripotency during naïve-to-primed conversion in mice. http://biorxiv.org/lookup/doi/10.1101/2020.09.25.313239.

Bowness JS, Nesterova TB, Wei G, Rodermund L, Almeida M, Coker H, Carter EJ, Kadaster A, Brockdorff N. 2022. Xist-mediated silencing requires additive functions of SPEN and Polycomb together with differentiation-dependent recruitment of SmcHD1. Cell Rep 39.

Cecalev D, Viçoso B, Galupa R. 2024. Compensation of gene dosage on the mammalian X. Development (Cambridge) 151.

Chen G, Schell JP, Benitez JA, Petropoulos S, Yilmaz M, Reinius B, Alekseenko Z, Shi L, Hedlund E, Lanner F, et al. 2016. Single-cell analyses of X Chromosome inactivation dynamics and pluripotency during differentiation. Genome Res 26: 1342–1354.

Cheng S, Pei Y, He L, Peng G, Reinius B, Tam PPL, Jing N, Deng Q. 2019. Single-Cell RNA-Seq Reveals Cellular Heterogeneity of Pluripotency Transition and X Chromosome Dynamics during Early Mouse Development. Cell Rep 26: 2593--2607.e3.

Deng Q, Ramsköld D, Reinius B, Sandberg R. 2014. Single-cell RNA-seq reveals dynamic, random monoallelic gene expression in mammalian cells. Science (1979) 343: 193–196.

Deng X, Berletch JB, Ma W, Nguyen DK, Hiatt JB, Noble WS, Shendure J, Disteche CM. 2013. Mammalian X upregulation is associated with enhanced transcription initiation, RNA half-life, and MOF-mediated H4K16 acetylation. Dev Cell 25: 55–68.

Deng X, Hiatt JB, Nguyen DK, Ercan S, Sturgill D, Hillier LW, Schlesinger F, Davis CA, Reinke VJ, Gingeras TR, et al. 2011. Evidence for compensatory upregulation of expressed X-linked genes in mammals, Caenorhabditis elegans and Drosophila melanogaster. Nat Genet 43: 1179–1185.

Di KN, Disteche CM. 2006. Dosage compensation of the active X chromosome in mammals. Nat Genet 38: 47–53.

Dobin A, Davis CA, Schlesinger F, Drenkow J, Zaleski C, Jha S, Batut P, Chaisson M, Gingeras TR. 2013. STAR: Ultrafast universal RNA-seq aligner. Bioinformatics 29: 15–21.

Gayen S, Maclary E, Hinten M, Kalantry S. 2016. Sex-specific silencing of X-linked genes by Xist RNA. Proc Natl Acad Sci U S A 113.

Gelbart ME, Larschan E, Peng S, Park PJ, Kuroda MI. 2009. Drosophila MSL complex globally acetylates H4K16 on the male X chromosome for dosage compensation. Nat Struct Mol Biol 16: 825–832.

Gontan C, Achame EM, Demmers J, Barakat TS, Rentmeester E, Van Ijcken W, Grootegoed JA, Gribnau J. 2012. RNF12 initiates X-chromosome inactivation by targeting REX1 for degradation. Nature 485: 386–390.

Grant J, Mahadevaiah SK, Khil P, Sangrithi MN, Royo H, Duckworth J, McCarrey JR, Vandeberg JL, Renfree MB, Taylor W, et al. 2012. Rsx is a metatherian RNA with Xist-like properties in X-chromosome inactivation. Nature 487: 254–258.

Graves JAM. 2016. Evolution of vertebrate sex chromosomes and dosage compensation. Nat Rev Genet 17: 33–46.

Gupta V, Parisi M, Sturgill D, Nuttall R, Doctolero M, Dudko OK, Malley JD, Eastman PS, Oliver B. 2006. Global analysis of X-chromosome dosage compensation. J Biol 5.

Huynh KO, Lee JT. 2003. Inheritance of a pre-inactivated paternal X chromosome in early mouse embryos. Nature 426: 857–862.

Jonkers I, Barakat TS, Achame EM, Monkhorst K, Kenter A, Rentmeester E, Grosveld F, Grootegoed JA, Gribnau J. 2009. RNF12 Is an X-Encoded Dose-Dependent Activator of X Chromosome Inactivation. Cell 139: 999–1011.

Julien P, Brawand D, Soumillon M, Necsulea A, Liechti A, Schütz F, Daish T, Grützner F, Kaessmann H. 2012. Mechanisms and evolutionary patterns of mammalian and avian dosage compensation. PLoS Biol.

Lahn BT, Page DC. 1999. Four evolutionary strata on the human X chromosome. Science (1979) 286: 964–967.

Langmead B, Salzberg SL. 2012. Fast gapped-read alignment with Bowtie 2. Nat Methods 9: 357–359. https://www.nature.com/articles/nmeth.1923 (Accessed July 3, 2025).

Lentini A, Cheng H, Noble JC, Papanicolaou N, Coucoravas C, Andrews N, Deng Q, Enge M, Reinius B. 2022. Elastic dosage compensation by X-chromosome upregulation. Nat Commun 13.

Li B, Dewey CN. 2011. RSEM: Accurate transcript quantification from RNA-Seq data with or without a reference genome. BMC Bioinformatics 12.

Lin H, Gupta V, Vermilyea MD, Falciani F, Lee JT, O’Neill LP, Turner BM. 2007. Dosage compensation in the mouse balances up-regulation and silencing of X-linked genes. PLoS Biol 5: 2809–2820.

Lin H, Halsall JA, Antczak P, O’Neill LP, Falciani F, Turner BM. 2011. Relative overexpression of X-linked genes in mouse embryonic stem cells is consistent with Ohno’s hypothesis. Nat Genet 43: 1169–1170.

Liu M, Yue Y, Chen X, Xian K, Dong C, Shi M, Xiong H, Tian K, Li Y, Zhang QC, et al. 2025. Genome-coverage single-cell histone modifications for embryo lineage tracing. Nature 640: 828–839.

Lyon MF. 1961. Gene Action in the X-chromosom of the mouse. Nature 190: 372–373.

Lyu Q, Yang Q, Hao J, Yue Y, Wang X, Tian J, An L. 2022. A small proportion of X-linked genes contribute to X chromosome upregulation in early embryos via BRD4-mediated transcriptional activation. Curr Biol. http://www.ncbi.nlm.nih.gov/pubmed/36108637 (Accessed October 16, 2022).

Mahadevaiah SK, Sangrithi MN, Hirota T, Turner JMA. 2020. A single-cell transcriptome atlas of marsupial embryogenesis and X inactivation. Nature 586: 612–617.

Majumdar S, Bammidi LS, Naik HC, Manhas A, Baro R, Kalita A, Naskar AJ, Nidharshan S, Bariha GS, Notani D, et al. 2025. Xist upstream deletion leads to dysregulation of Xist and autosomal gene expression. Genome Res 35: 1992–2010. https://pubmed.ncbi.nlm.nih.gov/40769712/ (Accessed October 6, 2025).

Mak W, Nesterova TB, De Napoles M, Appanah R, Yamanaka S, Otte AP, Brockdorff N. 2004. Reactivation of the Paternal X Chromosome in Early Mouse Embryos. Science (1979) 303: 666–669. https://www.science.org/doi/10.1126/science.1092674 (Accessed June 6, 2024).

Moindrot B, Cerase A, Coker H, Masui O, Grijzenhout A, Pintacuda G, Schermelleh L, Nesterova TB, Brockdorff N. 2015. A Pooled shRNA Screen Identifies Rbm15, Spen, and Wtap as Factors Required for Xist RNA-Mediated Silencing. Cell Rep 12: 562–572.

Monkhorst K, Jonkers I, Rentmeester E, Grosveld F, Gribnau J. 2008. X Inactivation Counting and Choice Is a Stochastic Process: Evidence for Involvement of an X-Linked Activator. Cell 132: 410–421.

Naik HC, Baro R, Sarkar A, Nayak M, Sunagar K, Gayen S. 2025. The role of m6A RNA methylation in the maintenance of X chromosome inactivation and X-to-autosome dosage compensation in early embryonic lineages. Stem Cell Reports.

Naik HC, Chandel D, Majumdar S, Arava M, Baro R, BV H, Hari K, Ayyamperumal P, Manhas A, Jolly MK, et al. 2024. Lineage-specific dynamics of loss of X upregulation during inactive-X reactivation. Stem Cell Reports 19: 1564–1582. http://www.cell.com/article/S2213671124002893/fulltext (Accessed November 22, 2024).

Naik HC, Hari K, Chandel D, Jolly MK, Gayen S. 2022. Single-cell analysis reveals X upregulation is not global in pre-gastrulation embryos. iScience 25.

Naik HC, Hari K, Chandel D, Mandal S, Jolly MK, Gayen S. 2021. Semicoordinated allelic-bursting shape dynamic random monoallelic expression in pregastrulation embryos. iScience 24: 102954. 10.1016/j.isci. (Accessed September 1, 2021).

Ohno S. 1967. Sex Chromosomes and Sex-linked Genes. Springer-Verlag, Berlin, Heidelberg, New York 68: 1375.

Okamoto I, Otte AP, Allis CD, Reinberg D, Heard E. 2004. Epigenetic Dynamics of Imprinted X Inactivation during Early Mouse Development. Science (1979) 303: 644–649. https://www.science.org/doi/10.1126/science.1092727 (Accessed June 6, 2024).

Pandey RS, Wilson Sayres MA, Azad RK. 2013. Detecting evolutionary strata on the human X chromosome in the absence of gametologous Y-linked sequences. Genome Biol Evol 5: 1863–1871.

Quinlan AR, Hall IM. 2010. BEDTools: A flexible suite of utilities for comparing genomic features. Bioinformatics 26: 841–842.

Ramos-Ibeas P, Sang F, Zhu Q, Tang WWC, Withey S, Klisch D, Wood L, Loose M, Surani MA, Alberio R. 2019. Pluripotency and X chromosome dynamics revealed in pig pre-gastrulating embryos by single cell analysis. Nat Commun 10.

Reinius B, Mold JE, Ramsköld D, Deng Q, Johnsson P, Michaëlsson J, Frisén J, Sandberg R. 2016. Analysis of allelic expression patterns in clonal somatic cells by single-cell RNA-seq. Nat Genet 48: 1430–1435.

Reinius B, Shi C, Hengshuo L, Sandhu KS, Radomska KJ, Rosen GD, Lu L, Kullander K, Williams RW, Jazin E. 2010. Female-biased expression of long non-coding RNAs in domains that escape X-inactivation in mouse. BMC Genomics 11.

Rodríguez-Delgado CL, Waters SA, Waters DP. 2014. Paternal X inactivation does not correlate with X chromosome evolutionary strata in marsupials. BMC Evol Biol 14.

Saiba R, Arava M, Gayen S. 2018. Dosage compensation in human pre-implantation embryos: X-chromosome inactivation or dampening? EMBO Rep 19: e46294.

Samanta MK, Gayen S, Harris C, Maclary E, Murata-Nakamura Y, Malcore RM, Porter RS, Garay PM, Vallianatos CN, Samollow PB, et al. 2022. Activation of Xist by an evolutionarily conserved function of KDM5C demethylase. Nat Commun 13: 2602. https://pubmed.ncbi.nlm.nih.gov/35545632/ (Accessed July 29, 2022).

Sandstedt SA, Tucker PK. 2004. Evolutionary Strata on the Mouse X Chromosome Correspond to Strata on the Human X Chromosome. Genome Res 14: 267. /pmc/articles/PMC327101/ (Accessed September 14, 2021).

Sangrithi MN, Royo H, Mahadevaiah SK, Ojarikre O, Bhaw L, Sesay A, Peters AHFM, Stadler M, Turner JMA. 2017. Non-Canonical and Sexually Dimorphic X Dosage Compensation States in the Mouse and Human Germline. Dev Cell 40: 289–301.e3.

Soh YQS, Alföldi J, Pyntikova T, Brown LG, Graves T, Minx PJ, Fulton RS, Kremitzki C, Koutseva N, Mueller JL, et al. 2014. Sequencing the mouse y chromosome reveals convergent gene acquisition and amplification on both sex chromosomes. Cell 159: 800–813.

Takagi N, Sasaki M. 1975. Preferential inactivation of the paternally derived X chromosome in the extraembryonic membranes of the mouse. Nature 1975 256:5519 256: 640–642. https://www.nature.com/articles/256640a0 (Accessed June 6, 2024).

Talon I, Janiszewski A, Theeuwes B, Lefevre T, Song J, Bervoets G, Vanheer L, De Geest N, Poovathingal S, Allsop R, et al. 2021. Enhanced chromatin accessibility contributes to X chromosome dosage compensation in mammals. Genome Biol 22.

Tong YB, Shi MW, Qian SH, Chen YJ, Luo ZH, Tu YX, Xiong YL, Geng YJ, Chen C, Chen ZX. 2021. GenOrigin: A comprehensive protein-coding gene origination database on the evolutionary timescale of life. Journal of Genetics and Genomics 48: 1122–1129.

Wang X, Douglas KC, VandeBerg JL, Clark AG, Samollow PB. 2014. Chromosome-wide profiling of X-chromosome inactivation and epigenetic states in fetal brain and placenta of the opossum, Monodelphis domestica. Genome Res 24: 70–83.

Wei C, Kesner B, Yin H, Lee JT. 2024. Imprinted X chromosome inactivation at the gamete-to-embryo transition. Mol Cell 84: 1442–1459.e7. https://pubmed.ncbi.nlm.nih.gov/38458200/ (Accessed December 15, 2025).

Xiao S, Xie D, Cao X, Yu P, Xing X, Chen CC, Musselman M, Xie M, West FD, Lewin HA, et al. 2012. Comparative epigenomic annotation of regulatory DNA. Cell 149: 1381–1392.

Yang F, Babak T, Shendure J, Disteche CM. 2010. Global survey of escape from X inactivation by RNA-sequencing in mouse. Genome Res 20: 614–622.

Yildirim E, Sadreyev RI, Pinter SF, Lee JT. 2012. X-chromosome hyperactivation in mammals via nonlinear relationships between chromatin states and transcription. Nat Struct Mol Biol 19: 56–62.

Zhang R, Yang M, Schreiber J, O’Day DR, Turner JMA, Shendure J, Noble WS, Disteche CM, Deng X. 2025. Cross-species imputation and comparison of single-cell transcriptomic profiles. Genome Biol 26.

Zhou F, Wang R, Yuan P, Ren Y, Mao Y, Li R, Lian Y, Li J, Wen L, Yan L, et al. 2019. Reconstituting the transcriptome and DNA methylome landscapes of human implantation. Nature 572: 660–664.

Zhu Q, Sang F, Withey S, Tang W, Dietmann S, Klisch D, Ramos-Ibeas P, Zhang H, Requena CE, Hajkova P, et al. 2021. Specification and epigenomic resetting of the pig germline exhibit conservation with the human lineage. Cell Rep 34.

Zou H, Yu D, Du X, Wang J, Chen L, Wang Y, Xu H, Zhao Y, Zhao S, Pang Y, et al. 2019. No imprinted XIST expression in pigs: biallelic XIST expression in early embryos and random X inactivation in placentas. Cellular and Molecular Life Sciences 76: 4525–4538.

