## Supplementary Figure for "Evolutionary origins and chromatin state shape X-chromosome upregulation pattern during eutherian and metatherian embryogenesis"

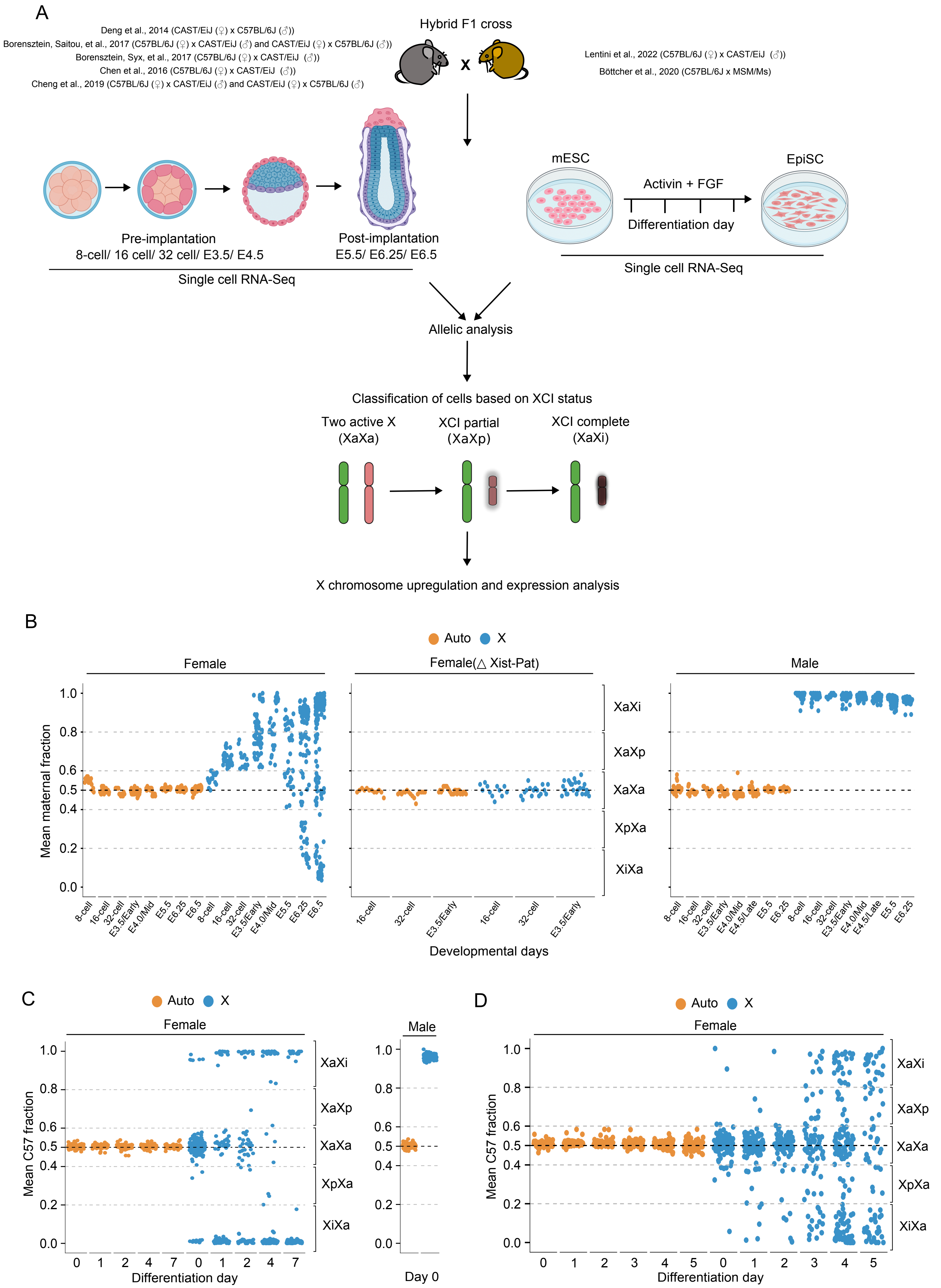
A
Deng et al., 2014 (CAST/EiJ (♀) x C57BL/6J (♂))
Borensztein, Saitou, et al., 2017 (C57BL/6J (♀) x CAST/EiJ (♂) and CAST/EiJ (♀) x C57BL/6J (♂))
Borensztein, Syx, et al., 2017 (C57BL/6J (♀) x CAST/EiJ (♂))
Chen et al., 2016 (C57BL/6J (♀) x CAST/EiJ (♂))
Cheng et al., 2019 (C57BL/6J (♀) x CAST/EiJ (♂) and CAST/EiJ (♀) x C57BL/6J (♂))
Hybrid F1 cross
x
Lentini et al., 2022 (C57BL/6J (♀) x CAST/EiJ (♂))
Böttcher et al., 2020 (C57BL/6J x MSM/Ms)
Pre-implantation
8-cell/ 16 cell/ 32 cell/ E3.5/ E4.5
Post-implantation
E5.5/ E6.25/ E6.5
Single cell RNA-Seq
mESC
Activin + FGF
Differentiation day
EpiSC
Single cell RNA-Seq
Allelic analysis
Classification of cells based on XCI status
Two active X
(XaXa)
XCI partial
(XaXp)
XCI complete
(XaXi)
X chromosome upregulation and expression analysis
B
Auto
X
Female
Female(△ Xist-Pat)
Male
Mean maternal fraction
1.0
0.8
0.6
0.5
0.4
0.2
0.0
XaXi
XaXp
XaXa
XpXa
XiXa
8-cell
16-cell
32-cell
E3.5/Early
E4.0/Mid
E4.5/Late
E5.5
E6.25
E6.5
Developmental days
C
Auto
X
Female
Male
Mean C57 fraction
Differentiation day
0
1
2
4
7
Day 0
XaXi
XaXp
XaXa
XpXa
XiXa
D
Auto
X
Female
Mean C57 fraction
Differentiation day
0
1
2
3
4
5
XaXi
XaXp
XaXa
XpXa
XiXa

**Fig. S1: Workflow and characterizing the XCI status of individual cells of mouse embryos and differentiated ESC (related to Fig. 1 to 6)**

(A) Schematic showing the analytical approach for characterizing the XCI status (cells with two active X: XaXa, cells with partially inactivated X: XaXi and cell with complete X-inactivation: XaXi) at the single cell level through allele-specific analysis of scRNA-seq datasets obtained from hybrid mouse embryos (pre- and post-implantation), ESC and differentiated ESC. Source of datasets and details of mouse strains were used to generate embryos and ESC have been provided. (B) Identification of XaXa, XaXp and XaXi cells in preimplantation embryos (WT and Xist-pat) based on fraction expression from maternal X-chromosome. As expected, expression of X-linked genes was solely from the maternal X in male cells (XmY). Autosomal genes showed equivalent expression from both maternal and paternal allele. (C) and (D) Identification of cells with different categories based on their XCI status (XaXa, XaXp and XaXi) during differentiation of ESC based on fraction expression from C57 allele. Male cells showed expression of X-linked genes from the C57 allele only as expected. Autosomal genes had equivalent expression from both alleles.

A

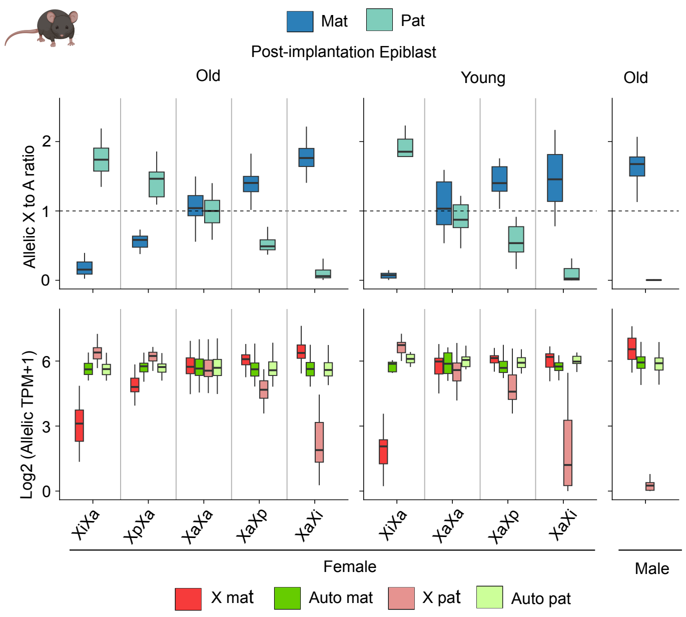

Mat
Pat
Post-implantation Epiblast
Old
Young
Old
Allelic X to A ratio
2
1
0
Log2 (Allelic TPM+1)
6
3
0
XiXa
XpXa
XaXa
XaXp
XaXi
XiXa
XaXa
XaXp
XaXi
Female
Male
X mat
Auto mat
X pat
Auto pat

**Fig. S2: Active-X upregulation analysis of old and young genes in post-implantation epiblast using the same set of genes used for analysis of pre-implantation embryos and post-implantation (ExE and VE) undergoing imprinted XCI (related to Fig. 2)**

Top: Plots representing the allelic X:A profile in XaXa, XaXp and XaXi cells at post-implantation epiblast cells; bottom: Allelic expression profile of X-linked and autosomal genes in old and young categories.

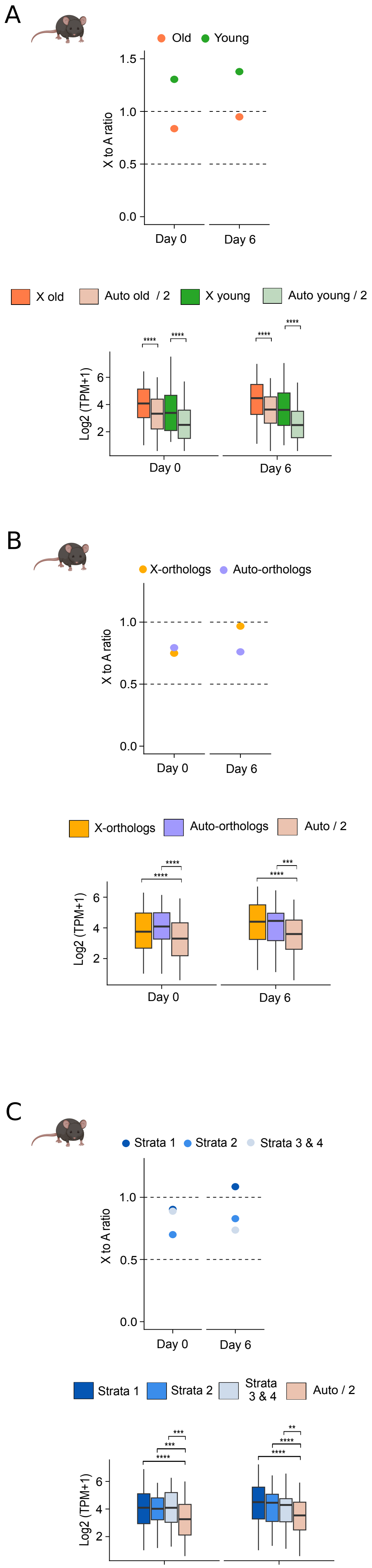
A
Old Young
X to A ratio
0.0
0.5
1.0
1.5
Day 0
Day 6
X old
Auto old / 2
X young
Auto young / 2
Log2 (TPM+1)
2
4
6
Day 0
Day 6
B
X-orthologs Auto-orthologs
X to A ratio
0.0
0.5
1.0
Day 0
Day 6
X-orthologs
Auto-orthologs
Auto / 2
Log2 (TPM+1)
2
4
6
Day 0
Day 6
C
Strata 1 Strata 2 Strata 3 & 4
X to A ratio
0.0
0.5
1.0
Day 0
Day 6
Strata 1
Strata 2
Strata 3 & 4
Auto / 2
Log2 (TPM+1)
2
4
6
Day 0
Day 6

**Fig. S3: X to A dosage compensation profiling in ESC and differentiated ESC (related to Fig. 7).**

X:A ratio (top) and expression plots (bottom) for (A) old and young (B) X-ortholog and auto-ortholog (C) strata1, strata2 and strata 3+4 genes in d0 and d6 differentiated ESC. Bulk RNA-seq datasets were used for this analysis. One-sided Wilcoxon rank-sum tests: P < 0.0001; ****, < 0.001; ***, < 0.01; **, < 0.05; *, NS; non-significant.
